# Implications of error-prone long-read whole-genome shotgun sequencing on characterizing reference microbiomes

**DOI:** 10.1101/2020.03.05.978866

**Authors:** Yu Hu, Li Fang, Christopher Nicholson, Kai Wang

## Abstract

Long-read sequencing techniques, such as the Oxford Nanopore Technology, can generate reads that are tens of kilobases in length, and are therefore particularly relevant for microbiome studies. However, due to the higher per-base error rates than typical short-read sequencing, the application of long-read sequencing on microbiomes remains largely unexplored. Here we deeply sequenced two human microbiota mock community samples (HM-276D and HM-277D) from the Human Microbiome Project. We showed that assembly programs consistently achieved high accuracy (~99%) and completeness (~99%) for bacterial strains with adequate coverage. We also found that long-read sequencing provides accurate estimates of species-level abundance (R=0.94 for 20 bacteria with abundance ranging from 0.005% to 64%). Our results demonstrate the feasibility to characterize complete microbial genomes and populations from error-prone Nanopore sequencing data, but also highlight necessary bioinformatics improvements for future metagenomics tool development.

## Background

The fundamental importance of microbiota as the microbial communities that reside in human body is increasingly recognized. Over the past decade, there have been tremendous amounts of evidence suggesting that microbiota plays a crucial role in human health through modulating the metabolic functions, as well as food energy harvest and storage. Microbiota, especially the gut microbiota, is associated with many chronic diseases such as obesity, diabetes, metabolic syndrome, inflammatory bowel disease (IBD), irritable bowel syndrome (IBS), liver disease, hepatocellular and colorectal carcinoma(Gill et al., 2006, Lewis et al., 2015, Chehoud et al., 2015, Hooper et al., 2003, Jones et al., 2015, Ley et al., 2006, Liang et al., 2015, Sartor, 2008, Schauber et al., 2003, Turnbaugh et al., 2009, Wang et al., 2016, Wen et al., 2008, Wu et al., 2011, Group et al., 2009). Therefore, accurate profiling of complete genomes and population are crucial to understanding the impact of microbiota on human health. Currently, high-throughput sequencing technologies have been widely used in microbial community characterization. In particular, 16S ribosomal RNA (rRNA)(Janda and Abbott, 2007) and shotgun metagenome sequencing on Illumina platforms(Quince et al., 2017) are two dominant approaches for describing microbiomes. Overall, the high-throughput nature of metagenomics sequencing allows us to interpret microbial community by using computational approaches such as operational taxonomic unit (OTU) identification(Hao and Chen, 2012), abundance quantification(Chen et al., 2017), read assembly(Ruan and Li, 2019, Bertrand et al., 2019, Koren et al., 2017, Kolmogorov et al., 2019, Li et al., 2015), binning and taxonomic profiling(Gregor et al., 2016, Huson et al., 2018, Huson et al., 2016, Francis et al., 2013, Hong et al., 2014, Byrd et al., 2014). Specifically, 16S rRNA sequencing targets on very specific regions that are highly variable between species, which is much cost-efficient. This is very useful for us to examine and compare the microbiota across high number of samples in a large scale project. However, this technique can only identify bacteria but not viruses or fungi, and the low resolution limits its usage in microbiome study below the genus level. As opposed to only the 16S sequences, shotgun metagenome sequencing surveys the whole genomes of all organism in the community (Jovel et al., 2016, Laudadio et al., 2018, Ranjan et al., 2016). It allows us to perform deep investigation of the microbial community as its ability to capture sequences from all organisms.

Despite the theoretical advantage of shotgun metagenome sequencing, due to the short read length (150 to 300 nucleotides), metagenomes cannot be fully characterized by next-generation sequencing (NGS) data. In addition, the lack of contextual information has become a barrier for short read to span both intra- and intergenomic repeats, which is crucial for complete de novo genome assembly of all dominant species in a microbial community. As a consequence, short-read assemblies remain highly fragmented. In comparison, the use of long-read sequencing has the potential to facilitate the complete and contiguous metagenome assembly. Lee *et al*. (Lee et al., 2014) sequenced a reference mock community sample using PacBio long read and evaluated the metagenome assembly performance. Results showed that single-molecule real-time (SMRT) long read data offered significantly improved assembly contiguity by spanning many of repetitive regions while single bacteria chromosome was assembled to more than 50 contigs based on short read data. In recent years, the Oxford Nanopore technologies (ONT) have offered advantages over traditional short-read NGS technologies in genome study. This single-molecule sequencing platform is able to generate average read length of >10kbp, spanning low complexity and repetitive genomic regions, which provides much more continuous assemblies. Subsequently, this approach has become an attractive option in metagenomics sequencing. While the ONT have great potential, complete and contiguous de novo metagenome assembly is still constrained by the high error rate (~15%) of single-molecule long-read sequence data(Sczyrba et al., 2017). Therefore, a comprehensive evaluation of long-read bioinformatics tools in microbial profiling is needed(Mason et al., 2017). Nicholls *et al*. (Nicholls et al., 2019) presented Nanopore sequencing data sets of two mock communities with 10 microbial species from ZymoBIOMICS(McIntyre et al., 2019). They showed the utility of these data sets for future bioinformatics method development for long-read metagenomics. However, publicly available data sets based other sequencing technologies of these samples are limited as the samples are only commercially available and are not well studied so far by competing approaches. A study to evaluate the advantages of Nanopore sequencing in complete microbial genomes and a comparison over other sequencing technologies is still lacking so far.

In this article, we generated two deeply sequenced Nanopore data sets from new reference samples that are more commonly studied, and performed comprehensive analysis to compare microbial community profiling performance with PacBio and Illumina technologies. We first generated 525× coverage data on HM-276D mock community sample from Human Microbiome Project, which is an evenly mixed DNA sample of 20 bacterial strains (each with 5% abundance). We performed de novo assembly analysis with 4 long-read assemblers at different depth of coverage. 20 bacterial genomes were assembled with high accuracy and genome completeness. This sample also has been well studied by many groups. As mentioned above, Lee et al. (Lee et al., 2014) sequenced this mock community with PacBio to show the improvement of long-read data in metagenome assembly analysis. Jones *et al*.(Jones et al., 2015) compared the influence of different NGS platforms on genomic and functional predictions using HM-276D sample. We downloaded these two data sets and compared the performance with Nanopore data. Our results show that Nanopore improved assembly contiguity compared to PacBio and Illumina across computational approaches. Next, we sequenced HM-277D Mock Community sample with 1068× coverage. HM-277D is unevenly mixed DNA sample of 20 bacterial strains. Kuleshove *et al*.(Kuleshov et al., 2016) sequenced this sample with Illumina TruSeq synthetic long read technique and showed the improvement in bacterial species identification, genome reconstruction compared to short sequences. Also, Leggett *et al*. (Leggett et al., 2018) demonstrated Nanopore metagenomics sequence can be reliably classified using this community. In addition to metagenome assembly, we evaluated taxonomy binning and profiling performance across technologies (Nanopore and PacBio) and samples (HM-276D and HM-277D). High identification and classification accuracy were achieved above the species level. Overall, we demonstrate the technical feasibility to characterize complete microbial genomes and populations from error-prone Nanopore sequencing without any DNA amplification. We also discuss the limitations of current bioinformatics tools, when dealing with error-prone long-read metagenomics sequencing data. All our data are made publicly available, to benefit computational tool development on long-read based microbial genome assembly for metagenomics studies.

## Results

### Sequence data quality

HM-276D DNA sample includes 20 evenly mixed bacteria strains with reference genome size 70 Mb in total with 39 chromosomes. 11,610,183 reads with 35,578,375,166 bases (525× coverage depth) were generated on the Nanopore GridION platform, with a median length of 1,374 bp. The N50 length is 6,828 bp and median read quality is 9.39 in Phred scale. By using minimap2, 95% of reads were successfully aligned to reference genomes of 20 bacterial strains with 13.1% error rate. As shown in **Figure 1A**, read coverage across 20 bacterial strains has good agreement with known abundances. Read depth is relatively homogenous across bacteria strains with 521.9X (sd = 524.7X) in average. Sequencing depth of each strain is at least 150 reads and only 0.03% region is covered by less than 3 reads.

**Figure 1.**
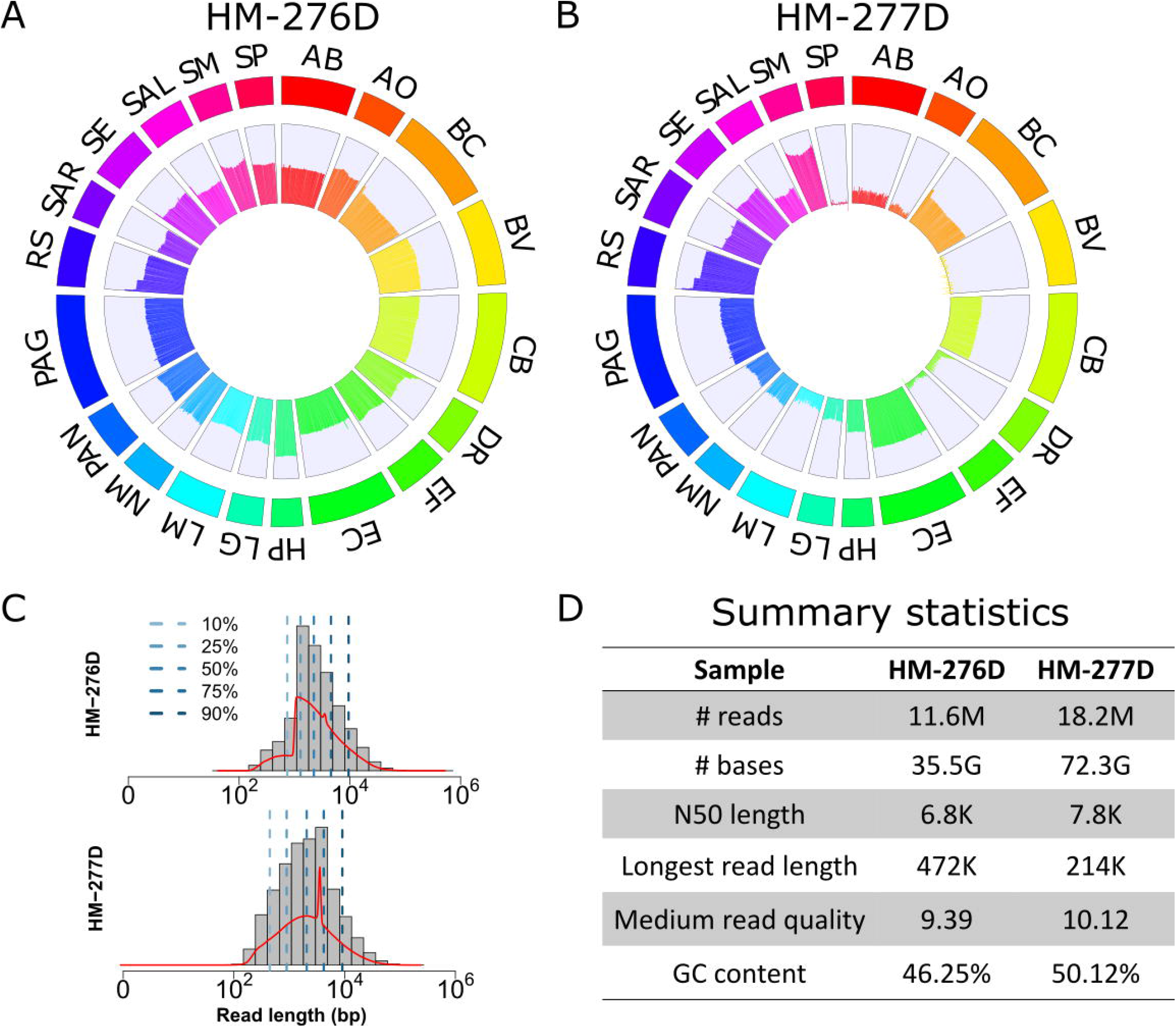
Summary of Nanopore Sequencing data from HM-276D and HM-277D microbial communities. **(A, B)** Circos plots of read coverage across whole genome of 20 bacterial strains from **(A)** HM-276D and **(B)** HM-277D. Each chromosome was divided to bins with 5,000 bp width. Average read coverage was calculated within each bin and converted to log scale to facilitate viewing and comparing between bacterial strains. AB, *Acinetobacter baumannii;* AO, *Actinomyces odontolyticus;* BC, *Bacillus cereus;* BV, *Bacteroides vulgatus;* CB, *Clostridium beijerinckii;* DR, *Deinococcus radiodurans;* DF, *Enterococcus faecalis;* EC, *Escherichia coli;* HP, *Helicobacter pylori;* LG, *Lactobacillus gasseri;* LM, *Listeria monocytogenes;* NM, *Neisseria meningitides;* PAN, *Propionibacterium acnes;* PAG, *Pseudomonas aeruginosa;* RS, *Rhodobacter sphaeroides;* SAR, *Staphylococcus aureus;* SE, *Staphylococcus epidermidis;* SAL, *Streptococcus agalactiae;* SM, *Streptococcus mutans;* SP, *Streptococcus pneumonia;* **(C)** Read length distribution of HM-276D and HM-277D data sets. Blue dashed lines represent different quantiles. Red line represents the density of read length distribution. **(D)** Summary statistics of HM-276D and HM-277D data sets. Each value was calculated by using pycoQC (Leger and Leonardi, 2019) and LongreadQC. Real-time statistics are shown in **Figure S1-S5**.

HM-277D DNA sample includes 20 unevenly mixed bacteria strains. 18,254,839 reads data set with 72,312,638,112 bases (1068× coverage depth) were generated, leading to 2,065 bp in median read length with 10.12 median read quality. The N50 length is 7,857 bp. 99.2% of QC-passed reads were mapped to the reference genome and the error rate was 9.8%. As shown in **Figure 1A**, read distribution is more heterogeneous across strains due to unevenly mixed samples. The average coverage is 988.8 reads with standard deviation =1941.6 bp. This leads to 1.6% of region with less than 3 reads covered and 4 strains with sequencing depth less than 10 bp, which makes it more difficult for biological interpretation of this microbial community.

### *De novo* assembly of HM-276D mock community

To assess the ability of Nanopore sequencing in profiling microbial community, we first conducted a de novo assembly of data set with 525× coverage from HM-276D mock community using 4 assemblers: wtdbg2(Ruan and Li, 2019), OPERA-MS(Bertrand et al., 2019), Canu(Koren et al., 2017) and meta-flye(Kolmogorov et al., 2019). OPERA-MS and meta-flye are designed to be capable of handling metagenome data, while wtdbg2 and canu are broadly used for haploid or diploid genomes. Overall, the results show promise for the characterization of microbial genomes using long-read sequencing data. Canu produced the largest assembly of 69.5 Mb (99.3% of the benchmark data), including 83 contigs with contig N50 length of 3.91 Mb. meta-flye assembled 67.7Mb genome with 89 contigs. wtdbg2 generated similar results with 64.9 Mb genome size, 61 contigs and 2.97 Mb N50 length. Assembly metrics of OPERA-MS (67.9 Mb genome size, 4734 contigs with contig N50 length of 2.94 Mb) are similar with Canu and wtdbg2 whereas much more contigs were generated because OPERA-MS utilizes both long and short sequencing reads for assembly. In addition, for aligned blocks, meta-flye yielded the highest NA50 with 1.71Mb in length compared to other assemblers (wtdbg2: 1.2Mb, OPERA-MS: 1.21Mb, Canu: 1.4Mb). Furthermore, by mapping all contigs to the reference genomes using MUMMer v3.23, we assessed the accuracy and genome completeness of contigs produced by 4 assemblers. As shown in **Figure 2A**, meta-flye achieved the highest genome fraction (99.99%) and 1-to-1 identity percentage (99.62%), followed by OPERA-MS (genome fraction: 99.98% and accuracy 99.92%), Canu (genome fraction 99.81% and accuracy 99.4%) and wtdbg2 (genome fraction 96.02% and accuracy 98.60%). Moreover, we evaluated aligned blocks for each method based on NA50 length. As shown in **Table S1**, meta-flye achieved the highest NA50 with 1.71Mb in length compared to other assemblers (wtdbg2: 1.2Mb, OPERA-MS: 1.21Mb, Canu: 1.4Mb). Overall, 4 tools generated results with similar good quality in term of contiguity, accuracy and completeness using long read data with evenly mixed samples at 525× coverage depth.

**Figure 2.**
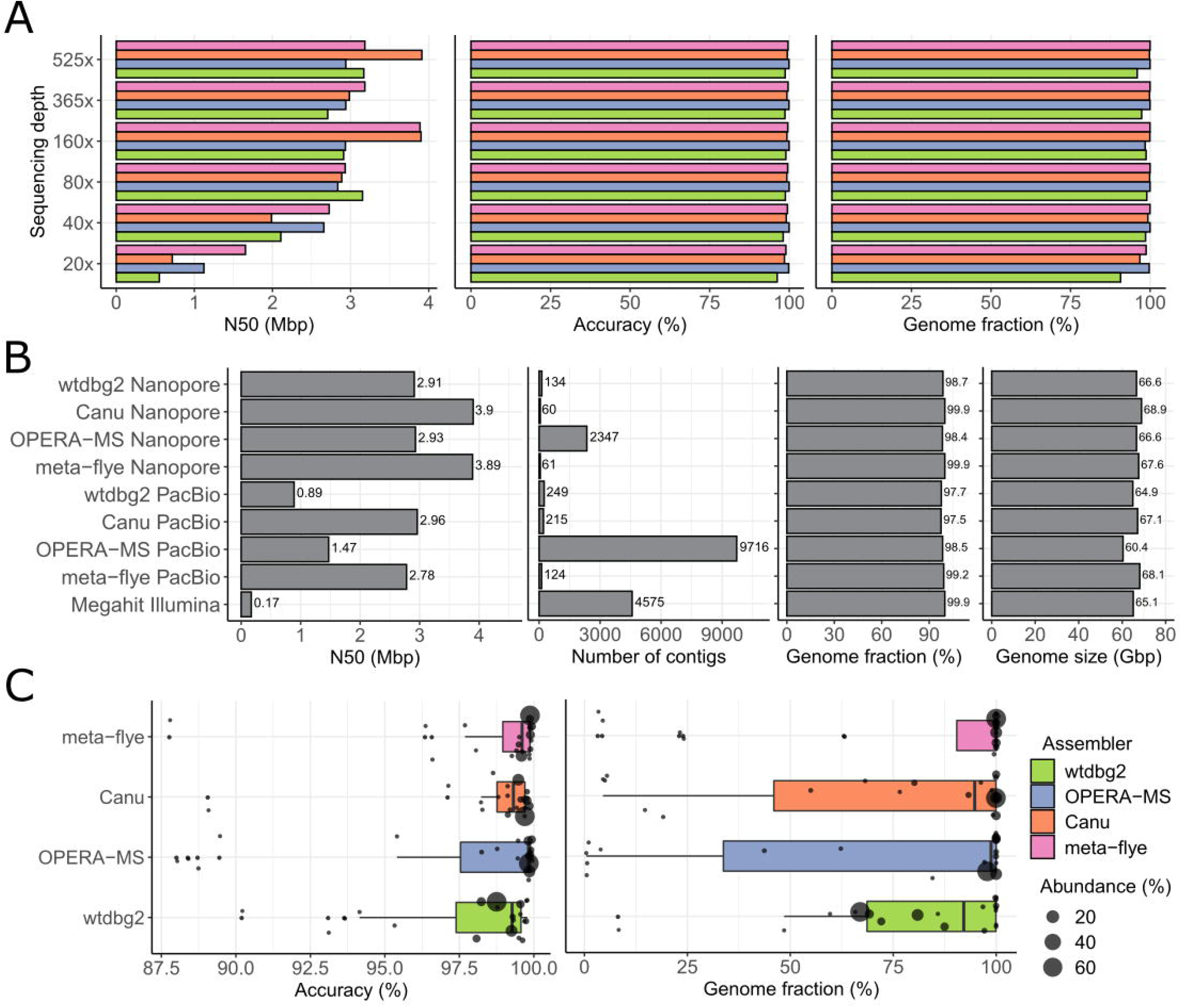
Assembly results for HM-276D and HM-277D data sets. **(A)** Assembly statistics (N50 length, accuracy and genome fraction) of each assembler at different coverage depths based on HM-276D data set. Colors indicate results from different assemblers (See **Supplemental material** for details in parameter settings). **(B)** Assembly statistics (N50 length, number of contigs, genome fraction and genome size) of each assembler based on HM-276D sample sequenced by different technologies (Nanopore, PacBio, Illumina). To make fair comparison, each data set was downsampled to 160× depth of coverage. **(C)** Strain-specific assembly performance of each assembler based on HM-277D data set. Assembly statistics (accuracy and genome fraction) distributions were presented using boxplots with jitter. Radius of each dot indicates the known relative abundance of each bacteria strain from the mock community.

Next, we subsampled 525× data set to 365× (70%), 160× (30%), 80× (15%), 40× (7.5%) and 20× (3.75%) to examine the effect of sequencing depths on de novo assembly (**Figure 2A**, **Table S1**). The assembly results of 4 tools ranges 95.95% to 99.96% in consensus accuracy and 91.26% to 99.99% in genome fraction. In specific, OPERA-MS outperforms others with the highest and most consistent metrics for completeness and accuracy across different sequencing depths because its metagenomics design substantially improves the robustness to low sequencing depth, where genome fractions are 99.68% in average (sd = 0.61%) and consensus identities are 99.92% in average (sd = 0.05%). Despite of reduced metrics as sequencing depth becoming lower, meta-flye and Canu still recovered at least 96.8% genomes with 98.5% accuracy. Notably, wtdbg2 improved the assembly metrics with coverage depth reduced from 365× to 80×. In addition, we examined whether genomes of 20 bacterial strains can be better constructed with Nanopore sequencing technology compared to PacBio and Illumina. As shown in **Figure 2B**, assemblers using Nanopore sequenced data outperforms other two technologies. With the same assembler, on average, the number of contigs of Nanopore is ~30% lower than PacBio, genome fraction and genome size are 1.56% and 3.1 Mb higher respectively. To understand the reason, we compared read length characteristics between these two data sets. For N50 length, ONT (7,350 bp) is 15% longer than PacBio (6,357 bp), and for N05 length, ONT (35.9 kbp) is even 159% longer than PacBio (13.8 kbp). This indicates that read length is the main advantage for ONT. Therefore, as shown in **Figure 2B**, N50 length of ONT (13.3 Mbp) is 68% longer than PacBio (8.1Mbp). Assemblies using Illumina sequenced data have 99.9% in accuracy, but more contigs generated and lower genome size in total compared to Nanopore.

### *De novo* assembly of HM-277D mock community

To evaluate the metagenome reconstruction in a more realistic setting, we carried out another de novo assembly of 1068× data set from HM-277D Mock Community, with unevenly mixed DNA samples of the 20 bacteria strains (**Figure S6**). Assembly accuracy still remains high, ranging from 97.78% to 99.75% across tools. However, not surprisingly, genome fractions and genome sizes of all methods are substantially lower than even community. This is because 13 bacterial strains have extremely low abundances (<1%) in this unevenly mixed samples, leading to reduced genome coverage fractions (Canu: 71.68%, OPERA-MS: 71.25%, meta-flye: 91.57%, wtdbg2: 82.95%) and genome sizes (Canu: 50.21 Mb, OPERA-MS: 47.99 Mb, meta-flye: 64.12 Mb, wtdbg2: 61.75 Mb). To assess how strain abundance affects assemblies, we calculated strain-specific genome fraction for each tool. Across bacterial strains, meta-flye recovered the highest percentage of genome (median 100%), followed by OPERA-MS (median: 98.75%), Canu (median 94.78%) and wtdbg2 (median: 91.66%) (**Figure 2C**). For bacteria with relative abundance higher than 0.2%, least 99.99% of reference genome can be covered by assembly contigs (meta-flye), with identity consensus reaching to 99.93%. These results suggest that bacterial strain with nontrivial abundance can be accurately assembled with Nanopore sequenced data. Overall, we observed that meta-flye returned assemblies for 20 bacterial strains with the best performance in completeness and accuracy. Metric for each strain is correlated with abundance of the corresponding bacteria. Some strains were proved hard to assemble for all assemblers due to extremely low relative abundance. For example, 13.6% of region of *Enterococcus faecalis* (0.011% relative abundance) were covered by 0 or 1 read and 56.1% covered by less than 3 reads, leading to 4.47% genome fraction for meta-flye. Moreover, there were 2 contigs belong to two different bacteria species, *Bacteroides vulgatus* (0.19% relative abundance) and *Streptococcus pneumoniae* (0.05% relative abundance), indicating the difficulty in differentiating one bacteria from another with low relative abundance.

### Taxon binning and identification

Metagenome assemblers construct contigs with variable length to recover original genome of each bacteria from microbial community. Subsequently, another major challenge in studying the identity and diversity of this community member is to classify sequenced reads or contigs correctly according to their taxonomic origins. Here we investigated the taxonomic binning performance based on 3 scenarios of long-read sequencing data, HM-276D (Nanopore, PacBio) and HM-277D (Nanopore) at 160× depth of coverage, using a state-of-art taxonomic binner Megan-LR. First, all long reads were aligned to NCBI-nr database. Then, we used Megan-LR with interval-union LCA algorithm to assign ~2 million aligned reads (~4.6 Mb bases) to taxonomic nodes (**Figure 3A,B, Figure S7-S10)**). Overall, 4.22 Mb (0.087%) from Nanopore data of HM-276D sample were mis-assigned, while 4.37 Mb (0.075%) and 4.66 Mb (0.141%) for Nanopore data of HM-277D and PacBio data of HM-276D respectively. Specifically, we evaluated the recovery of taxon bins at different ranks. We considered two metrics to quantify the read assignment accuracy, average precision and sensitivity of 20 bacteria strains. For each taxonomic bin, we obtained precision by calculating the percentage of reads correctly classified out of all binned reads. Sensitivity is the percentage of correctly assigned reads out of all reads originally from the bin. As shown in **Figure 3C**, HM-276D (Nanopore) has the highest precision, which are all above 60% from phylum to genus. HM-277D (Nanopore) followed, with all above 50%, while HM-276D (PacBio) has the lowest average precision due to predicted small false positive bins at the species level. Sensitivity has similar pattern (**Figure 3D**). HM-276D (Nanopore) still appears to the best data set for read classification than other two and the difference in accuracy between these 3 scenarios is similar across ranks. Nanopore is ~8% higher than PacBio and HM-276D is 10% higher than HM-277D. To evaluate the stability of read assignment accuracy, we calculated 95% confidence interval of precision and sensitivity for each scenarios at each rank. Not surprisingly, confidence bands are narrower at higher rank, indicating that more taxon recovery accuracy can be reached.

**Figure 3.**
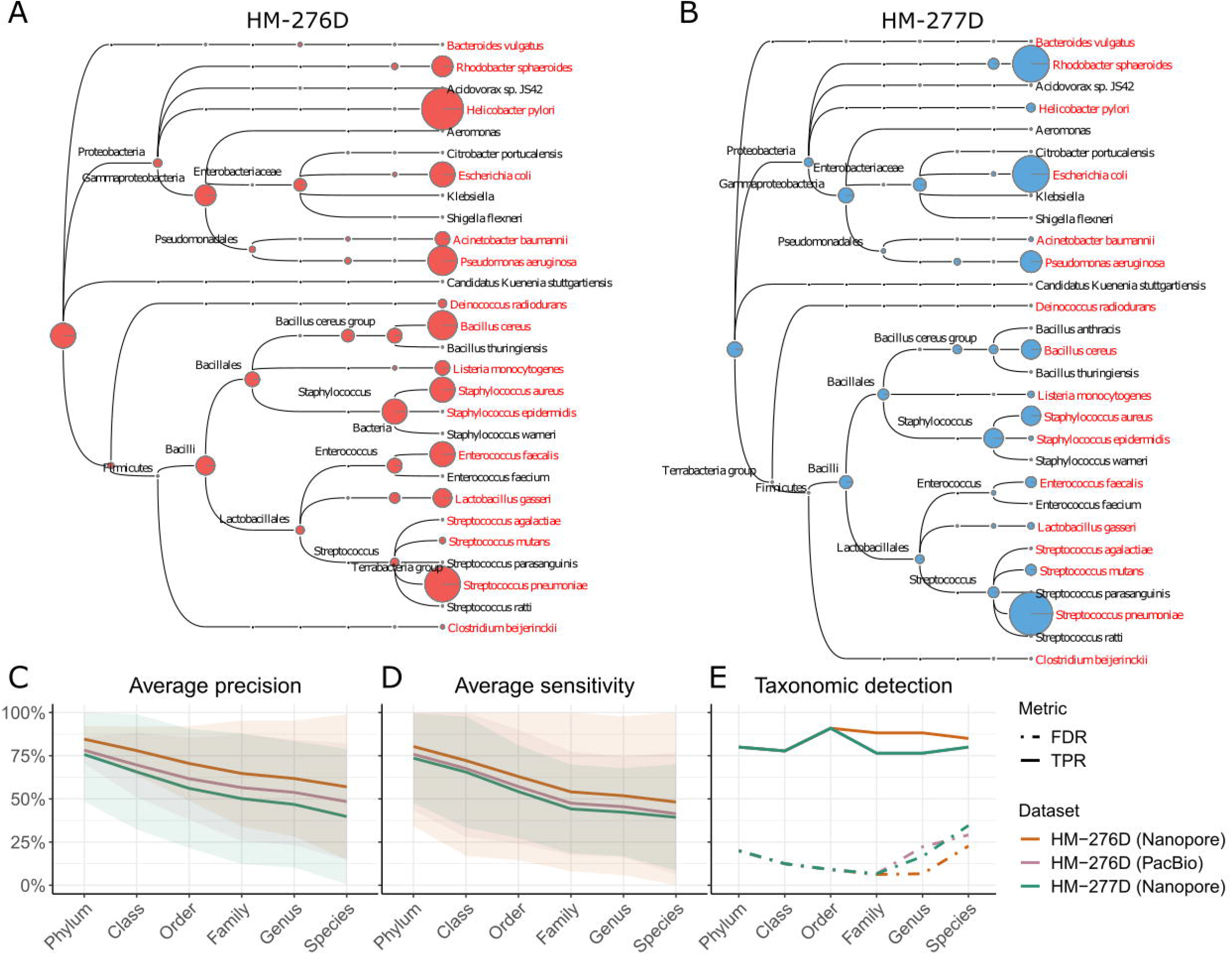
Taxonomic binning results for HM-276D and HM-277D data sets. **(A,B)** Megan taxonomic tree assignment obtained from HM-276D **(A)** and HM-277D **(B)** Nanopore sequenced data sets. Both data sets were downsampled to 160× depth of coverage. Each read was aligned against NCBI-nr protein reference data base, then binned and visualized using Megan-LR. Megan taxonomic tree showing bacteria taxa identified and their corresponding abundances across taxonomic rank. The radius of circle represents the number of reads assigned for each taxa. Bacterial strains highlighted in red represent true organisms in the mock community. **(C-E)** Taxonomic binning and identification performance metrics across ranks based on different data sets (indicated by colors). Average precision **(C)**, average sensitivity **(D)** and their 95% CIs were calculated based on metrics from different taxon at each rank. **(E)** Taxonomic detection accuracy metrics, true positive rate (solid) and false positive rate (dashed), were calculated based on identified taxon (reads > 10) at each rank. To make fair comparison, each data set was downsampled to 160× depth of coverage.

Owing to unevenly mixed bacteria strains, sensitivity is much more variable for HM-277D than other HM-276D. Overall, these results demonstrated the advantage of long-read data in accurate taxon recovery above the family level, while binning accuracy and stability were relatively at the species level.

In addition to assigning sequence fragments (reads or contigs) to taxon bins, we recognized the importance of accurate determination of taxonomic identity presence or absence from microbial community. Therefore, we continued to investigate the performance of taxonomic identity prediction between data from HM-276D (Nanopore, PacBio) and HM-277D (Nanopore). For taxon prediction, we defined that the species is significantly present in the community when at least 10 reads were assigned to it, while identity with less 10 supporting reads was marked as absence. We considered two other metrics to quantify the detection accuracy, true positive rate (TPR) and false discover rate (FDR), where TPR is the percentage of correctly predicted taxonomic identities out of known existing taxon and FDR is the percentage of incorrectly predicted taxonomic identities out of all predicted taxon. TPR and FDR were calculated at different ranks in **Figure 3E**. TPR were consistent across 3 data sets from phylum to order level (90%-77%). Below the order level, PacBio (HM-276D) and Nanopore (HM-277D) are 22% lower compared to Nanopore (HM-276D) (92%-87%). From phylum to family level, FDRs were controlled under 15% for all 3 data sets. However, at the genus level, more than 20% of detections are false for PacBio (HM-276D) and Nanopore (HM-277D) while 6% for Nanopore (HM-276). All 3 scenarios have inflated FDR (>20%) at the species level. Across data sets, there was drastic increase in FDR between phylum to family level and below family level, 10%±3% and 21%±5%. Similar to binning results, Nanopore data of HM-276D still consistently performed better than other two data sets across ranks. However, accurately predicting taxonomic profiles at the species level still remains challenging due to many false predicted taxonomic identities with 10 to 100 reads assigned incorrectly.

### Strain profiling

Despite the challenges in assembly and binning of HM-277D microbial community even at the species level, especially for low abundance bacteria (relative abundance < 1%), the golden standard profile of this mock community still allows us to evaluate other unique advantages of this deeply sequenced data set at strain level. First, we examined the ability in identifying these 13 extremely rare strains based on annotated target genes. To explore the sensitivity of strain detection using this data set, we mapped raw sequenced reads to reference genomes of the 20 bacterial strains with Minimap2. Then, for each strain-specific gene, the average coverage were estimated by summing up read depth across all exonic region, normalized for gene length. In addition, exon coverage fractions were calculated. We required a gene with average coverage greater than 1 and exon coverage fraction greater 50% simultaneously in order to be declared as a detected gene. The results are shown in **Figure 4A** and **Figure S11-S12**. Detection rates and average coverage among all genes largely keep high in abundant strains (>1%), ranging from 96.4 bp to 4207.6 bp, as well as most of rare strains (<1%). Most of bacterial strains except for *Bacteroides vulgatus* (69.1%) and *Streptococcus pneumoniae* (81.7%) have achieved at least 97% gene detection rate.

**Figure 4.**
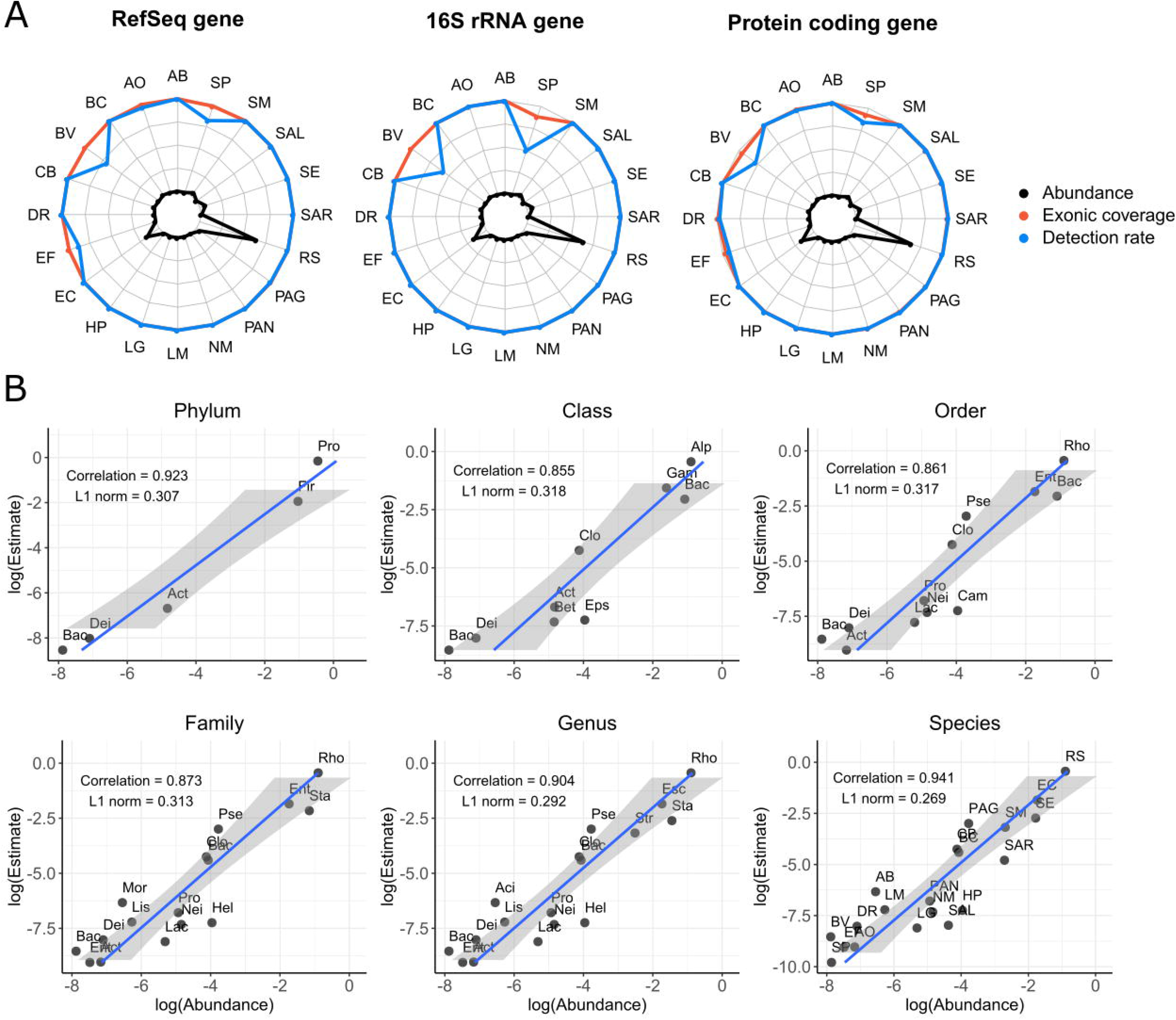
Taxonomic profiling results for HM-277D data sets. **(A)** Gene identification performance of 20 bacterial strains. 3 gene sets (RefSeq, 16S rRNA, protein coding) were evaluated. Colors indicate different metrics (exonic coverage and detection rate). Exonic coverage (orange) is the percentage of exonic region covered by at least 1 read out of all exons. Detection rate (blue) is the percentage of genes with coverage depth > 1 and exonic coverage > 50% out of all genes. Gold standard abundance of each strain was indicated in black. **(B)** Bacteria abundance estimation. Scatter plots abundance estimates versus gold standard abundances from HM-277D mock community across taxonomic ranks. Abundances were converted to log scale to facilitate viewing. Pearson correlation and L1 norm were utilized to quantify the performance. Estimates consistently share a good agreement with gold standard across ranks with correlation > 0.85 and L1 norm < 0.32. Abbreviations for bacterial name above the species level are listed below. Phylum level: Actinobacteria, Bacteroidetes (Bac), Deinococcus-Thermus (Dei), Firmicutes (Fir), Proteobacteria (Pro); Class level: Actinobacteria (Act), Alphaproteobacteria (Alp), Bacilli (Bac), Bacteroidia (Bact), Betaproteobacteria (Bet), Clostridiales (Clo), Deinococcus (Dei), Epsilonproteobacteria (Eps), Gammaproteobacteria (Gam); Order level: Actinomycetales (Act), Bacillales (Bac), Bacteroidales (Bact), Campylobacterales (Cam), Clostridiales (Clo), Deinococcales (Dei), Enterobacteriales (Ent), Lactobacillales (Lac), Neisseriaceae (Nei), Propionibacteriaceae (Pro), Pseudomonadales (Pse), Rhodobacterales (Rho); Family level: Actinomycetaceae (Act), Bacillaceae (Bac), Bacteroidaceae (Bact), Clostridiaceae (Clo), Deinococcaceae (Dei), Enterobacteriaceae (Ent), Enterococcaceae (Ent), Helicobacteraceae (Hel), Lactobacillaceae (Lac), Listeriaceae (Lis), Moraxellaceae (Mor), Neisseriaceae (Nei), Propionibacteriaceae (Pro), Pseudomonadaceae (Pse), Rhodobacteraceae (Rho), Staphylococcaceae (Sta); Genus level: Acinetobacter (Act), Actinomyces (Act), Bacillus (Bac), Bacteroides (Bact), Clostridium (Clo), Deinococcus (Dei), Enterococcus (Ent), Escherichia (Esc), Helicobacter (Hel), Lactobacillus (Lac), Listeria (Lis), Neisseria (Nei), Propionibacterium (Pro), Pseudomonas (Pse), Rhodobacter (Rho), Staphylococcus (Sta), Streptococcus (Str).

Next, we recognized that 16S rRNA genes are most commonly used as gene marker for bacteria identification, we further selected them out for each strain based RefSeq annotation. As shown in **Figure 4A**, though *Bacteroides vulgatus* and *Streptococcus pneumoniae* still have about 50% of 16S rRNA genes undetected by raw sequenced reads, 18 strains have 100% detection rates and exon coverage fraction with 434.77 bp coverage in average, which demonstrates the feasibility of identifying rare strain (<1%) in microbial community with long-read sequencing data. Additionally, read coverage of protein coding genes for 20 bacterial strains was summarized, which shows similar results. 14 strains have average coverage above 100 bp and gene detection rates for 18 strains have reached to 99%, indicating the presence of bacterial strains in the sample.

To understand the composition, diversity and spatial dynamics of microbial communities, we continued to evaluate the bacterial abundance estimation accuracy based on Nanopore data. We determined two abundance metrics to measure the accuracy, Pearson correlation and L1 norm. These two metrics assess how well Nanopore sequenced reads can reconstruct the bacterial abundances in comparison to the gold standard. Relative abundance was obtained by normalizing total read coverage with chromosome length for each taxon at different ranks. As shown in **Figure 4B**, abundance estimates at the species level agrees well with the known relative abundances from the mock community. However, abundance estimation at higher ranks appears to be more challenging, as correlation coefficient ranges from 0.87 to 0.85 and L1 norm is above 0.3 from class to family level, while two metrics improved with Pearson correlation > 0.9 and L1 < 0.29 when rank is below the family level. Poor abundance estimation at class or family level may due to the presence of extremely rare bacterial strains in the HM-277D sample, as read coverages were simply summed up between species belonging to the same family or class without accounting for abundance heterogeneity.

## Discussion

Complete genome assembly and population profiling are critical for the interpretation of microbial community diversity. However, a benchmarking long-read data set with consistent evaluation metrics is still lacking, which has hindered our understanding of long-read sequence data in metagenome assembly. In this study, we deeply sequenced HM-276D and HM-277D samples to assess the performance of error-prone Nanopore sequencing data and bioinformatics tools in characterizing microbial community. Assemblers consistently achieved high accuracy and completeness for nontrivial bacteria strains and genome binners performed well at above the genus level. Furthermore, by targeting on marker genes, we were able to identify rare strains with extremely low abundance in microbial community. Overall, our results have demonstrated that the technical feasibility to characterize complete microbial genomes and populations from Nanopore sequencing data with metagenomic software.

We note that despite the feasibility to characterize complete microbial genomes from long-read sequencing data, there are still challenges to be resolved in our study. Even for evenly mixed samples, the best performing assembler meta-flye achieve 99.99% consensus accuracy. However, as the reference genomes contains 70 Mb, 0.04% error rate has led to 28 Kbp of mismatches. These erroneous bases could be due to sequencing errors in low quality read, a major drawback of long-read sequence data and base modification, which may complicate the genome assembly. To prevent these errors, a sequencer with unbiased and methylation-aware base caller is in need. (We also acknowledge that some of the mismatches may be due to natural differences between reference microbiome samples and the reference genomes that were used.) In addition, there is still room for further improvement in assembly completeness by using longer reads or better designed assemblers to account for long repeats in genomes. In our study, we assembled long-read sequenced data from 20 bacterial strains across species. However, the performance at strain-level still remains unknown as closely related genomes is always a major challenge for genome assembly. In the future, we anticipate that more mock microbial community will be released with bacteria at strain level for benchmarking study.

By evaluating the performance of bioinformatics tools across different technologies, we found that third generation sequencing generally facilitates the complete characterization of complex bacterial genomes by overcoming many limitations of second generation sequencing. The short read length has limited the ability of Illumina sequencing in genome interpretation. For example, the length of repetitive genomic region is larger than a single read. As a consequence, intra- and intergenomic diversities are unlikely to be captured by short sequencing data. This issue has been resolved by long-read sequencing technologies (ONT and PacBio), which is able to span low complexity and repetitive regions by providing sequence reads with at least 10 kb in length. While generating data with much higher error rate than PacBio, ONT has become a promising platform in many applications, especially for studies requiring large amounts of data. This is because ONT provides longer reads (up to 900 kb in length) with higher throughput compared to PacBio (10-15 kb in length). Moreover, ONT is currently more affordable with lower per-base cost of data generation, which is a key factor in long-read sequencing studies. Overall, the application of these two major long-read sequencing platforms in metagenomics analysis of complex communities is still restricted by higher error rate. This problem could be addressed with improvement of consensus sequences. Recently, newly released R10 chip from ONT has longer base-contacting constriction in the pore, which improves the homopolymer resolution as compared to R9 and improved per-base error rates. Similarly, the HiFi protocol from PacBio can provide Sanger-quality accuracy (>99%) with reduced read length, which are still much longer than short-read sequencing for assembly of complex genomes (Wenger et al., 2019).This can lead to metagenome assembly with higher accuracy and completeness, as well as more accurate OTU identification. Future metagenomics studies are expected to be changed dramatically by this approach. For example, strain UA159 and NN2025 under species *Streptococcus mutans* only share 8% common regions, which can be uniquely assigned. We then found that 20% of ONT reads can cover the unique region of these two strains respectively, which is infeasible for short reads. Therefore, with better quality of long-read data, this approach may allow us to identify bacteria of interest directly at strain level instead of performing binning analysis in the future.

In addition to illustrating the advantages brought by long-read sequence data, we also assessed the performance of four *de novo* assembly algorithms and a long-read genome binner. The bioinformatics challenges to interpret rich information from complex microbial community include high error rates and low throughput for long-read sequencing, fragmented nature for short-read sequencing, and large CPU hours requirement. For evenly mixed (each with 5% abundance) HM-276D mock community, 4 tools consistently achieved high accuracy and completeness. No single assembler significantly outperforms others. By subsampling data to less coverage depths, not surprisingly, we found that the corresponding metrics for 4 tools decreased. In terms of speed, wtdbg2 is tens of times faster than other tools. For the unevenly mixed mock community HM-277D, assembly accuracy still remain high for all 4 tools (~97-98%). Genome fraction was reduced because 13 rare bacterial strains (<1%) were poorly assembled. Hybrid-assembler OPERA-MS, which combines the advantages from long and short-read technologies, shows more robust performance to bacterial strains with extremely low abundance than other tools. However, it produced much more contigs with less contiguity while meta-flye, Canu and wtdbg2 returned single contig for 18, 15 and 17 strains respectively. Furthermore, taxonomic binning results show that Megan-LR performs well when genomes are not closely related. Taxon bins were reconstructed with acceptable accuracy down to the genus level while performance decreased at species and strain level.

In summary, our results demonstrate the feasibility to characterize complete microbial genomes and populations from error-prone Nanopore sequencing data, but also highlight necessary bioinformatics improvements for future metagenomics tool development to handle specific challenges in error-prone long-read sequencing data. We believe that future metagenomics studies will benefit from this approach to assemble complete microbial genomes, while maintaining the theoretical ability to detect DNA methylations and base modifications, infer repetitive elements and structural variants, and achieve strain-level resolution within microbial communities. All the data sets on reference microbiomes are made publicly available to facilitate benchmarking studies on metagenomics and the development of novel software tools.

## Limitations of the Study

In this study, we note that there is still room for further improvement in assembly completeness using long reads. Also, the performance of binning analysis using long-read at strain-level still remains unknown.

## Resource Availability

### Lead Contact

Further information and requests for resources and reagents should be directed to and will be fulfilled by the Lead Contact, Kai Wang

### Materials Availability

This study did not generate new unique reagents

**Table 1.** Mapping statistics of HM-276D and HM-277D sequenced data set. Sequenced data were mapped against reference genomes of 20 known bacterial strains. Sequences indicates the number of QC passed reads. Number of mapped and unmapped reads were summarized. MQ0 represents number of mapped reads with MQ=0.Clipping was ignored when calculating total length, bases mapped. Bases mapped (cigar) provides a more accurate number of mapped bases. Number of mismatches were obtained from NM field of BAM file.

| **Mapping statistics** | **HM-276D** | **HM-277D** |
| --- | --- | --- |
| **# of reads** | 8,086,684 | 18,254,839 |
| **# of mapped reads** | 7,640,934 | 18,110,317 |
| **reads unmapped** | 445,750 | 144,522 |
| **reads MQ0** | 60,972 | 103,601 |
| **non-primary alignments** | 287,369 | 732,671 |
| **total length** | 33,563,573,383 | 72,312,638,112 |
| **bases mapped** | 32,143,689,158 | 72,216,146,980 |
| **bases mapped (cigar)** | 31,156,025,998 | 70,073,211,829 |
| **mismatches** | 4,104,593,752 | 6,925,222,080 |
| **average length** | 4,150 | 3,961 |
| **maximum length** | 472,762 | 214,792 |
| **average Phred quality per base** | 13 | 17 |

## Data and Code Availability

The Oxford Nanopore sequencing data that support the findings of this study have been deposited in the BioProject database at http://www.ncbi.nlm.nih.gov/bioproject/630658 (reference number: **PRJNA630658**)

The PacBio data used in this study was generated from the PacBio RS II sequencer. The data was downloaded from the following URL: https://github.com/PacificBiosciences/DevNet/wiki/Human_Microbiome_Project_MockB_Shotgun.

The Illumina paired-end data for HM-276D was downloaded from NCBI SRA database with accession numbers **SRR2726671** and **SRR2726672**.

The Illumina TruSeq synthetic long-read data for HM-277D was downloaded from NCBI SRA database with accession number **SRR2822457**.

## Acknowledgements

This work was supported by CHOP Research Institute and NIH/NIGMS grant GM132713 to K.W..

The following reagent was obtained through BEI Resources, NIAID, NIH as part of the Human Microbiome Project: Genomic DNA from Microbial Mock Community B (Staggered, High Concentration), v5.2H, for Whole Genome Shotgun Sequencing, HM-277D.

The following reagent was obtained through BEI Resources, NIAID, NIH as part of the Human Microbiome Project: Genomic DNA from Microbial Mock Community B (Even, High Concentration), v5.1H, for Whole Genome Shotgun Sequencing, HM-276D.

## Author Contributions

Y.H. performed data analysis and wrote the manuscript. L.F designed the study, performed long-read sequencing and analyzed the data. C.N. performed long-read sequencing. K.W. designed the study, supervised the study and wrote the manuscript. All authors read, revised and approved the manuscript.

## References

Bertrand, D., Shaw, J., Kalathiyappan, M., Ng, A. H. Q., Kumar, M. S., Li, C., Dvornicic, M., Soldo, J. P., Koh, J. Y., Tong, C., Ng, O. T., Barkham, T., Young, B., Marimuthu, K., Chng, K. R., Sikic, M. & Nagarajan, N. 2019. Hybrid metagenomic assembly enables high-resolution analysis of resistance determinants and mobile elements in human microbiomes. Nat Biotechnol, 37, 937–944.

Byrd, A. L., Perez-Rogers, J. F., Manimaran, S., Castro-Nallar, E., Toma, I., Mccaffrey, T., Siegel, M., Benson, G., Crandall, K. A. & Johnson, W. E. 2014. Clinical PathoScope: rapid alignment and filtration for accurate pathogen identification in clinical samples using unassembled sequencing data. BMC Bioinformatics, 15, 262.

Chehoud, C., Albenberg, L. G., Judge, C., Hoffmann, C., Grunberg, S., Bittinger, K., Baldassano, R. N., Lewis, J. D., Bushman, F. D. & Wu, G. D. 2015. Fungal Signature in the Gut Microbiota of Pediatric Patients With Inflammatory Bowel Disease. Inflamm Bowel Dis, 21, 194856.

Chen, E. Z., Bushman, F. D. & Li, H. 2017. A Model-Based Approach For Species Abundance Quantification Based On Shotgun Metagenomic Data. Stat Biosci, 9, 13–27.

Francis, O. E., Bendall, M., Manimaran, S., Hong, C., Clement, N. L., Castro-Nallar, E., Snell, Q., Schaalje, G. B., Clement, M. J., Crandall, K. A. & Johnson, W. E. 2013. Pathoscope: species identification and strain attribution with unassembled sequencing data. Genome Res, 23, 1721–9.

Gill, S. R., Pop, M., Deboy, R. T., Eckburg, P. B., Turnbaugh, P. J., Samuel, B. S., Gordon, J. I., Relman, D. A., Fraser-Liggett, C. M. & Nelson, K. E. 2006. Metagenomic analysis of the human distal gut microbiome. Science, 312, 1355–9.

Gregor, I., Droge, J., Schirmer, M., Quince, C. & Mchardy, A. C. 2016. PhyloPythiaS+: a self-training method for the rapid reconstruction of low-ranking taxonomic bins from metagenomes. PeerJ, 4, el603.

Group, N. H. W., Peterson, J., Garges, S., Giovanni, M., Mcinnes, P., Wang, L., Schloss, J. A., Bonazzi, V., Mcewen, J. E., Wetterstrand, K. A., Deal, C., Baker, C. C., Dl Francesco, V., Howcroft, T. K., Karp, R. W., Lunsford, R. D., Wellington, C. R., Belachew, T., Wright, M., Giblin, C., David, H., Mills, M., Salomon, R., Mullins, C., Akolkar, B., Begg, L., Davis, C., Grandison, L., Humble, M., Khalsa, J., Little, A. R., Peavy, H., Pontzer, C., Portnoy, M., Sayre, M. H., Starke-Reed, P., Zakhari, S., Read, J., Watson, B. & Guyer, M. 2009. The NIH Human Microbiome Project. Genome Res, 19, 2317–23.

Hao, X. & Chen, T. 2012. OTU analysis using metagenomic shotgun sequencing data. PLoS One, 7, e49785.

Hong, C., Manimaran, S., Shen, Y., Perez-Rogers, J. F., Byrd, A. L., Castro-Nallar, E., Crandall, K. A. & Johnson, W. E. 2014. PathoScope 2.0: a complete computational framework for strain identification in environmental or clinical sequencing samples. Microbiome, 2, 33.

Hooper, L. V., Stappenbeck, T. S., Hong, C. V. & Gordon, J. I. 2003. Angiogenins: a new class of microbicidal proteins involved in innate immunity. Nat Immunol, 4, 269–73.

Huson, D. H., Albrecht, B., Bagci, C., Bessarab, I., Gorska, A., Jolic, D. & Williams, R. B. H. 2018. MEGAN-LR: new algorithms allow accurate binning and easy interactive exploration of metagenomic long reads and contigs. Biol Direct, 13, 6.

Huson, D. H., Beier, S., Flade, I., Gorska, A., El-Hadidi, M., Mitra, S., Ruscheweyh, H. J. & Tappu, R. 2016. MEGAN Community Edition - Interactive Exploration and Analysis of Large-Scale Microbiome Sequencing Data. PLoS Comput Biol, 12, e1004957.

Janda, J. M. & Abbott, S. L. 2007. 16S rRNA gene sequencing for bacterial identification in the diagnostic laboratory: pluses, perils, and pitfalls. J Clin Microbiol, 45, 2761–4.

Jones, M. B., Highlander, S. K., Anderson, E. L., Li, W., Dayrit, M., Klitgord, N., Fabani, M. M., Seguritan, V., Green, J., Pride, D. T., Yooseph, S., Biggs, W., Nelson, K. E. & Venter, J. C. 2015. Library preparation methodology can influence genomic and functional predictions in human microbiome research. Proc Natl Acad Sci USA, 112, 14024–9.

Jovel, J., Patterson, J., Wang, W., Hotte, N., O’Keefe, S., Mitchel, T., Perry, T., Kao, D., Mason, A. L., Madsen, K. L. & Wong, G. K. 2016. Characterization of the Gut Microbiome Using 16S or Shotgun Metagenomics. Front Microbiol, 7, 459.

Kolmogorov, M., Yuan, J., Lin, Y. & Pevzner, P. A. 2019. Assembly of long, error-prone reads using repeat graphs. Nat Biotechnol, 37, 540–546.

Koren, S., Walenz, B. P., Berlin, K., Miller, J. R., Bergman, N. H. & Phillippy, A. M. 2017. Canu: scalable and accurate long-read assembly via adaptive k-mer weighting and repeat separation. Genome Res, 27, 722–736.

Kuleshov, V., Jiang, C., Zhou, W., Jahanbani, F., Batzoglou, S. & Snyder, M. 2016. Synthetic long-read sequencing reveals intraspecies diversity in the human microbiome. Nat Biotechnol, 34, 649.

Laudadio, I., Fulci, V., Palone, F., Stronati, L., Cucchiara, S. & Carissimi, C. 2018. Quantitative Assessment of Shotgun Metagenomics and 16S rDNA Amplicon Sequencing in the Study of Human Gut Microbiome. OMICS, 22, 248–254.

Lee, C. H., Bowman, B. & Hall, R. Developments in PacBio^®^ metagenome sequencing: Shotgun whole genomes and full-length 16S. International Plant and Animal Genome Conference Asia, 2014.

Leger, A. & Leonardi, T. 2019. pycoQC, interactive quality control for Oxford Nanopore Sequencing. The Journal of Open Source Software, 4, 1236.

Leggett, R. M., Alcon-Giner, C., Heavens, D., Caim, S., Brook, T. C., Kujawska, M., Martin, S., Hoyles, L., Clarke, P. & Hall, L. J. 2018. Rapid profiling of the preterm infant gut microbiota using nanopore sequencing aids pathogen diagnostics. bioRxiv, 180406.

Lewis, J. D., Chen, E. Z., Baldassano, R. N., Otley, A. R., Griffiths, A. M., Lee, D., Bittinger, K., Bailey, A., Friedman, E. S., Hoffmann, C., Albenberg, L., Sinha, R., Compher, C., Gilroy, E., Nessel, L., Grant, A., Chehoud, C., Li, H., Wu, G. D. & Bushman, F. D. 2015. Inflammation, Antibiotics, and Diet as Environmental Stressors of the Gut Microbiome in Pediatric Crohn’s Disease. Cell Host Microbe, 18, 489–500.

Ley, R. E., Turnbaugh, P. J., Klein, S. & Gordon, J. I. 2006. Microbial ecology: human gut microbes associated with obesity. Nature, 444, 1022–3.

Li, D., Liu, C. M., Luo, R., Sadakane, K. & Lam, T. W. 2015. MEGAHIT: an ultra-fast single-node solution for large and complex metagenomics assembly via succinct de Bruijn graph. Bioinformatics, 31, 1674–6.

Liang, X., Bittinger, K., Li, X., Abernethy, D. R., Bushman, F. D. & Fitzgerald, G. A. 2015. Bidirectional interactions between indomethacin and the murine intestinal microbiota. Elife, 4, e08973.

Mason, C. E., Afshinnekoo, E., Tighe, S., Wu, S. & Levy, S. 2017. International Standards for Genomes, Transcriptomes, and Metagenomes. J Biomol Tech, 28, 8–18.

Mcintyre, A. B. R., Alexander, N., Grigorev, K., Bezdan, D., Sichtig, H., Chiu, C. Y. & Mason, C. E. 2019. Single-molecule sequencing detection of N6-methyladenine in microbial reference materials. Nat Commun, 10, 579.

Nicholls, S. M., Quick, J. C., Tang, S. & Loman, N. J. 2019. Ultra-deep, long-read nanopore sequencing of mock microbial community standards. Gigascience, 8.

Quince, C., Walker, A. W., Simpson, J. T., Loman, N. J. & Segata, N. 2017. Shotgun metagenomics, from sampling to analysis. Nat Biotechnol, 35, 833–844.

Ranjan, R., Rani, A., Metwally, A., Mcgee, H. S. & Perkins, D. L. 2016. Analysis of the microbiome: Advantages of whole genome shotgun versus 16S amplicon sequencing. Biochem Biophys Res Commun, 469, 967–77.

Ruan, J. & Li, H. 2019. Fast and accurate long-read assembly with wtdbg2. Nat Methods.

Sartor, R. B. 2008. Microbial influences in inflammatory bowel diseases. Gastroenterology, 134, 57794.

Schauber, J., Svanholm, C., Termen, S., Iffland, K., Menzel, T., Scheppach, W., Melcher, R., Agerberth, B., Luhrs, H. & Gudmundsson, G. H. 2003. Expression of the cathelicidin LL-37 is modulated by short chain fatty acids in colonocytes: relevance of signalling pathways. Gut, 52, 735–41.

Sczyrba, A., Hofmann, P., Belmann, P., Koslicki, D., Janssen, S., Droge, J., Gregor, I., Majda, S., Fiedler, J., Dahms, E., Bremges, A., Fritz, A., Garrido-Oter, R., Jorgensen, T. S., Shapiro, N., Blood, P. D., Gurevich, A., Bai, Y., Turaev, D., Demaere, M. Z., Chikhi, R., Nagarajan, N., Quince, C., Meyer, F., Balvociute, M., Hansen, L. H., Sorensen, S. J., Chia, B. K. H., Denis, B., Froula, J. L., Wang, Z., Egan, R., Don Kang, D., Cook, J. J., Deltel, C., Beckstette, M., Lemaitre, C., Peterlongo, P., Rizk, G., Lavenier, D., Wu, Y. W., Singer, S. W., Jain, C., Strous, M., Klingenberg, H., Meinicke, P., Barton, M. D., Lingner, T., Lin, H. H., Liao, Y. C., Silva, G. G. Z., Cuevas, D. A., Edwards, R. A., Saha, S., Piro, V. C., Renard, B. Y., Pop, M., Klenk, H. P., Goker, M., Kyrpides, N. C., Woyke, T., Vorholt, J. A., Schulze-Lefert, P., Rubin, E. M., Darling, A. E., Rattei, T. & Mchardy, A. C. 2017. Critical Assessment of Metagenome Interpretation-a benchmark of metagenomics software. Nat Methods, 14, 1063–1071.

Turnbaugh, P. J., Hamady, M., Yatsunenko, T., Cantarel, B. L., Duncan, A., Ley, R. E., Sogin, M. L., Jones, W. J., Roe, B. A., Affourtit, J. P., Egholm, M., Henrissat, B., Heath, A. C., Knight, R. & Gordon, J. I. 2009. A core gut microbiome in obese and lean twins. Nature, 457, 480–4.

Wang, F., Kaplan, J. L., Gold, B. D., Bhasin, M. K., Ward, N. L., Kellermayer, R., Kirschner, B. S., Heyman, M. B., Dowd, S. E., Cox, S. B., Dogan, H., Steven, B., Ferry, G. D., Cohen, S. A., Baldassano, R. N., Moran, C. J., Garnett, E. A., Drake, L., Otu, H. H., Mirny, L. A., Libermann, T. A., Winter, H. S. & Korolev, K. S. 2016. Detecting Microbial Dysbiosis Associated with Pediatric Crohn Disease Despite the High Variability of the Gut Microbiota. Cell Rep, 14, 945–955.

Wen, L., Ley, R. E., Volchkov, P. Y., Stranges, P. B., Avanesyan, L., Stonebraker, A. C., Hu, C., Wong, F. S., Szot, G. L., Bluestone, J. A., Gordon, J. I. & Chervonsky, A. V. 2008. Innate immunity and intestinal microbiota in the development of Type 1 diabetes. Nature, 455, 110913.

Wenger, A. M., Peluso, P., Rowell, W. J., Chang, P. C., Hall, R. J., Concepcion, G. T., Ebler, J., Fungtammasan, A., Kolesnikov, A., Olson, N. D., Topfer, A., Alonge, M., Mahmoud, M., Qian, Y., Chin, C. S., Phillippy, A. M., Schatz, M. C., Myers, G., Depristo, M. A., Ruan, J., Marschall, T., Sedlazeck, F. J., Zook, J. M., Li, H., Koren, S., Carroll, A., Rank, D. R. & Hunkapiller, M. W. 2019. Accurate circular consensus long-read sequencing improves variant detection and assembly of a human genome. Nat Biotechnol, 37, 1155–1162.

Wu, G. D., Chen, J., Hoffmann, C., Bittinger, K., Chen, Y. Y., Keilbaugh, S. A., Bewtra, M., Knights, D., Walters, W. A., Knight, R., Sinha, R., Gilroy, E., Gupta, K., Baldassano, R., Nessel, L., Li, H., Bushman, F. D. & Lewis, J. D. 2011. Linking long-term dietary patterns with gut microbial enterotypes. Science, 334, 105–8.

