## Supplemental material for "Implications of error-prone long-read whole-genome shotgun sequencing on characterizing reference microbiomes"

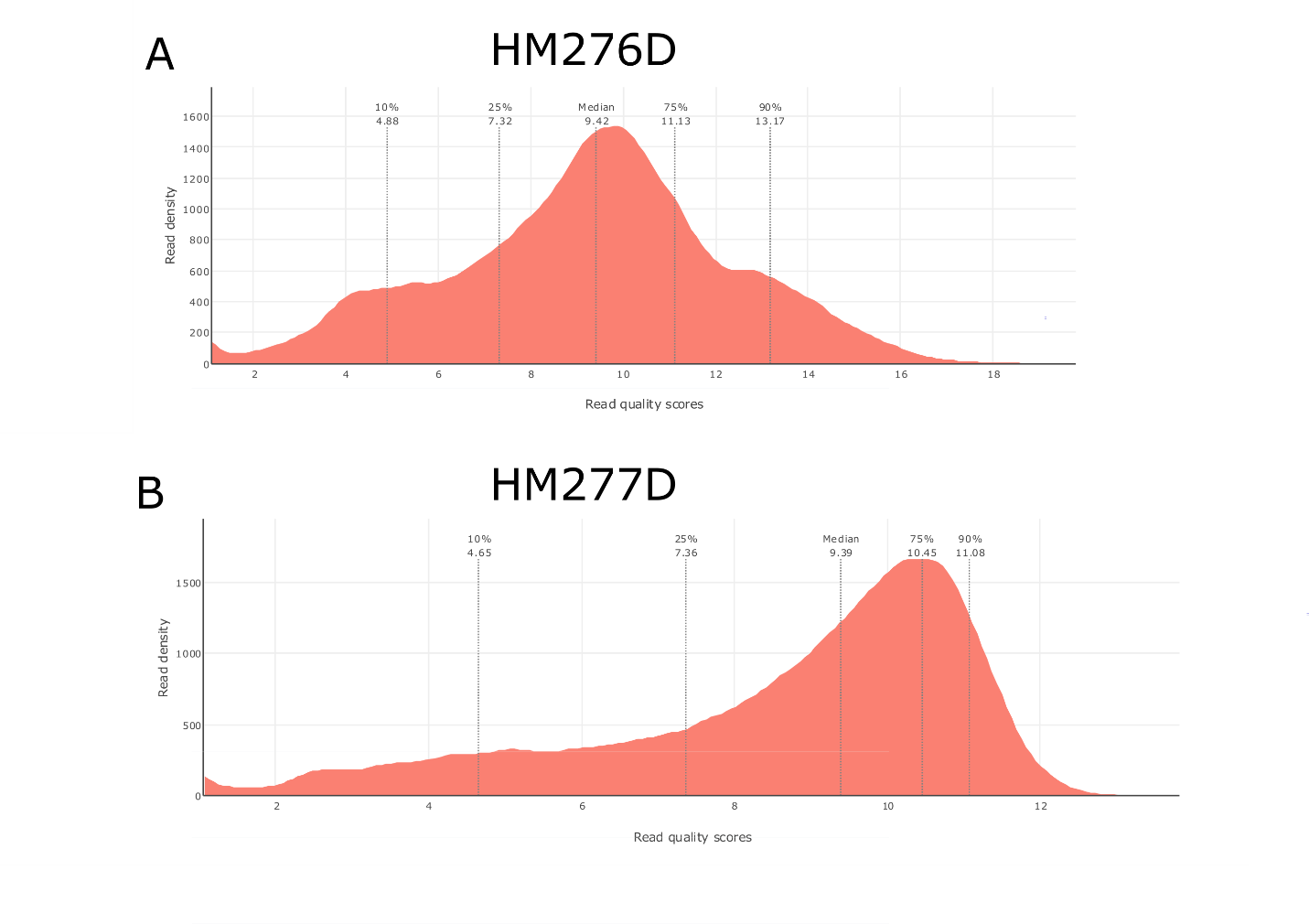

**Figure S1. Read quality of Nanopore sequencing data.** Read quality of sequenced data sets, HM-276D **(A)** and HM-277D **(B)**, were summarized using PycoQC respectively. Dashed lines indicate different quantiles (10%, 25%, 50%, 75%, 90%).

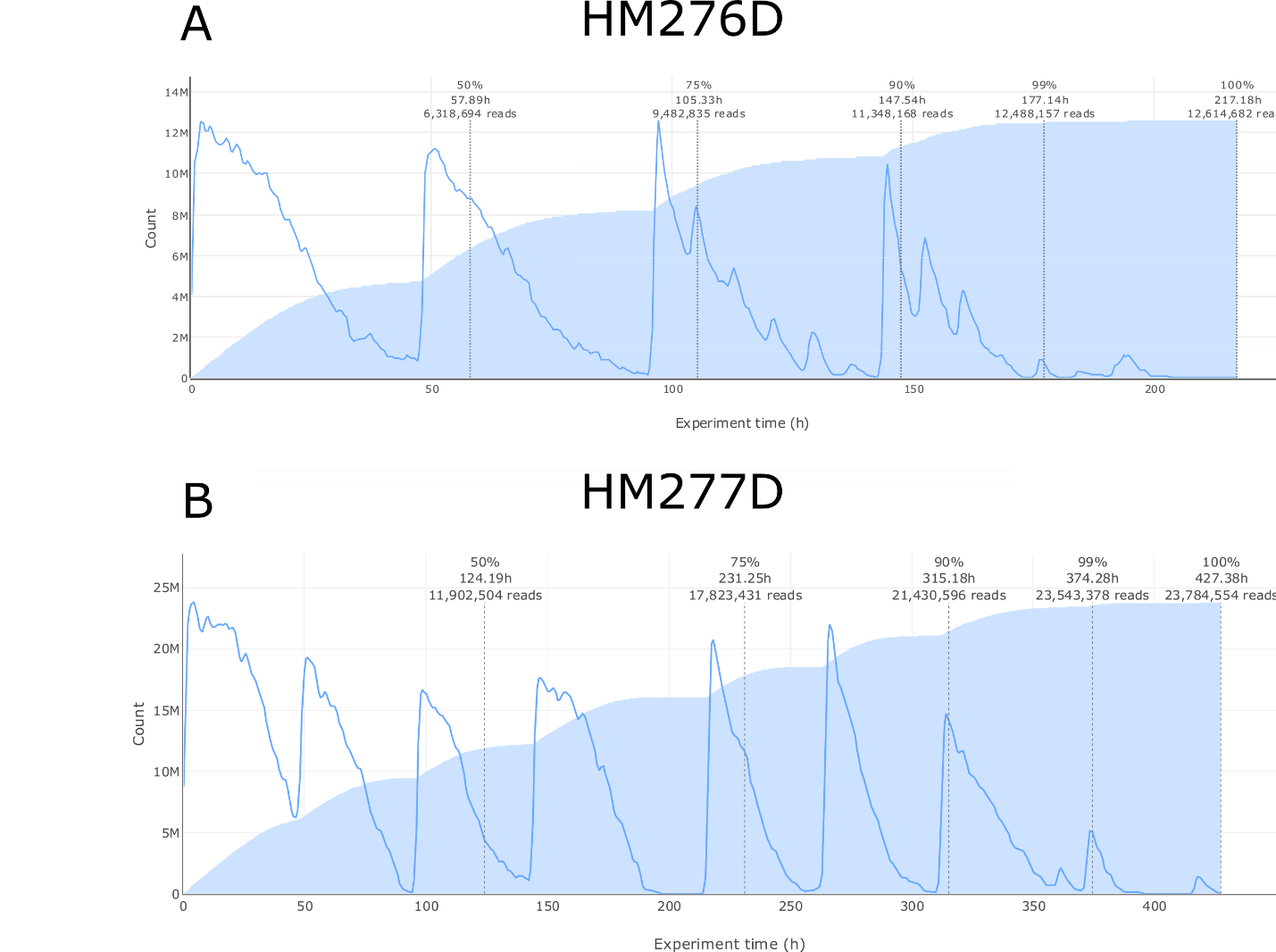

**Figure S2. Read output over experiment of Nanopore sequencing data.** Number of output reads over experiment time for sequenced data sets, HM-276D **(A)** and HM-277D **(B)**, were summarized using PycoQC. Blue line indicates output velocity at specific time. Shaded area represents cumulative read output over experiment time.

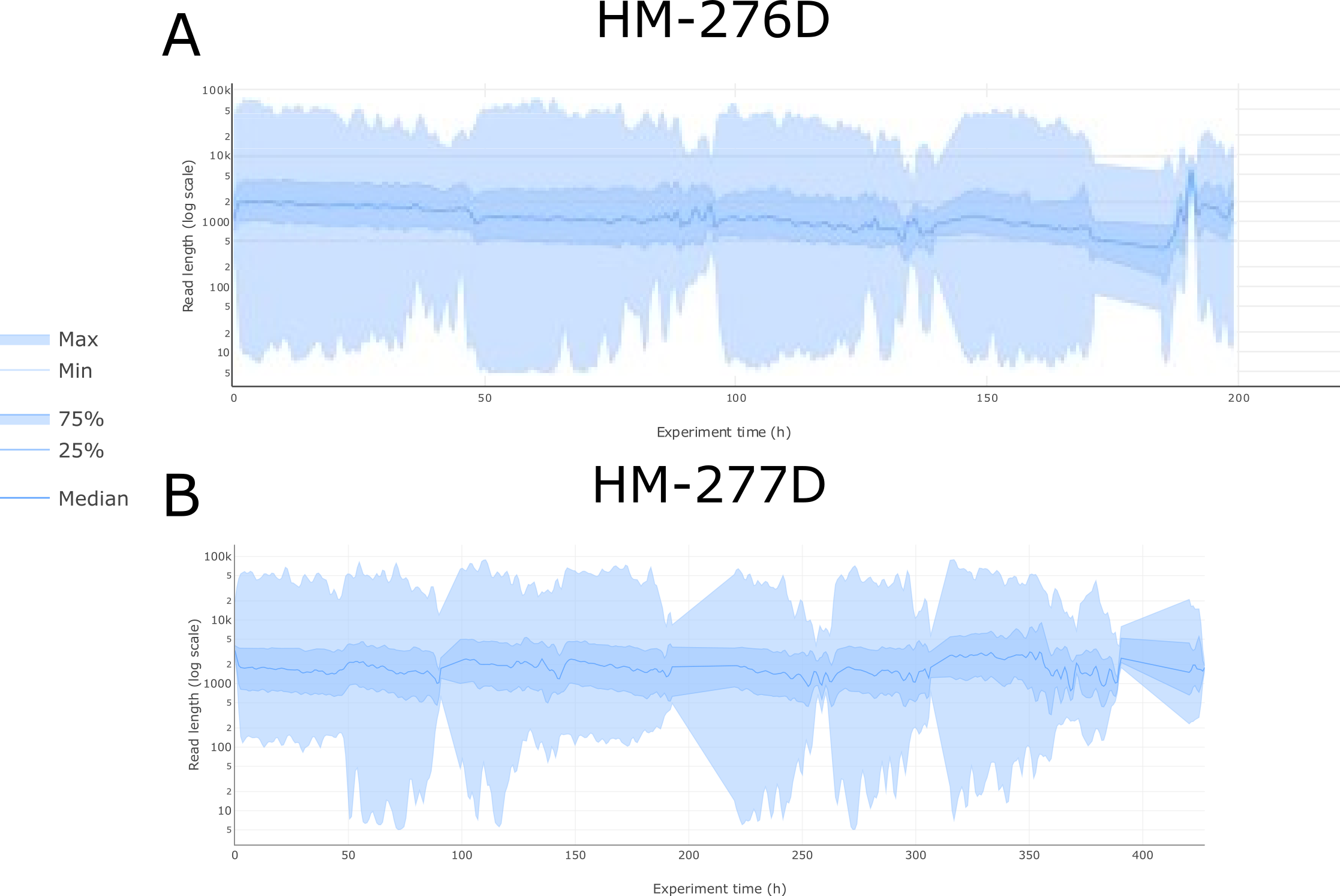

**Figure S3. Read length over experiment of Nanopore sequencing data.** Read length in log scale over experiment time for sequenced data sets, HM-276D **(A)** and HM-277D **(B)**, were summarized using PycoQC.

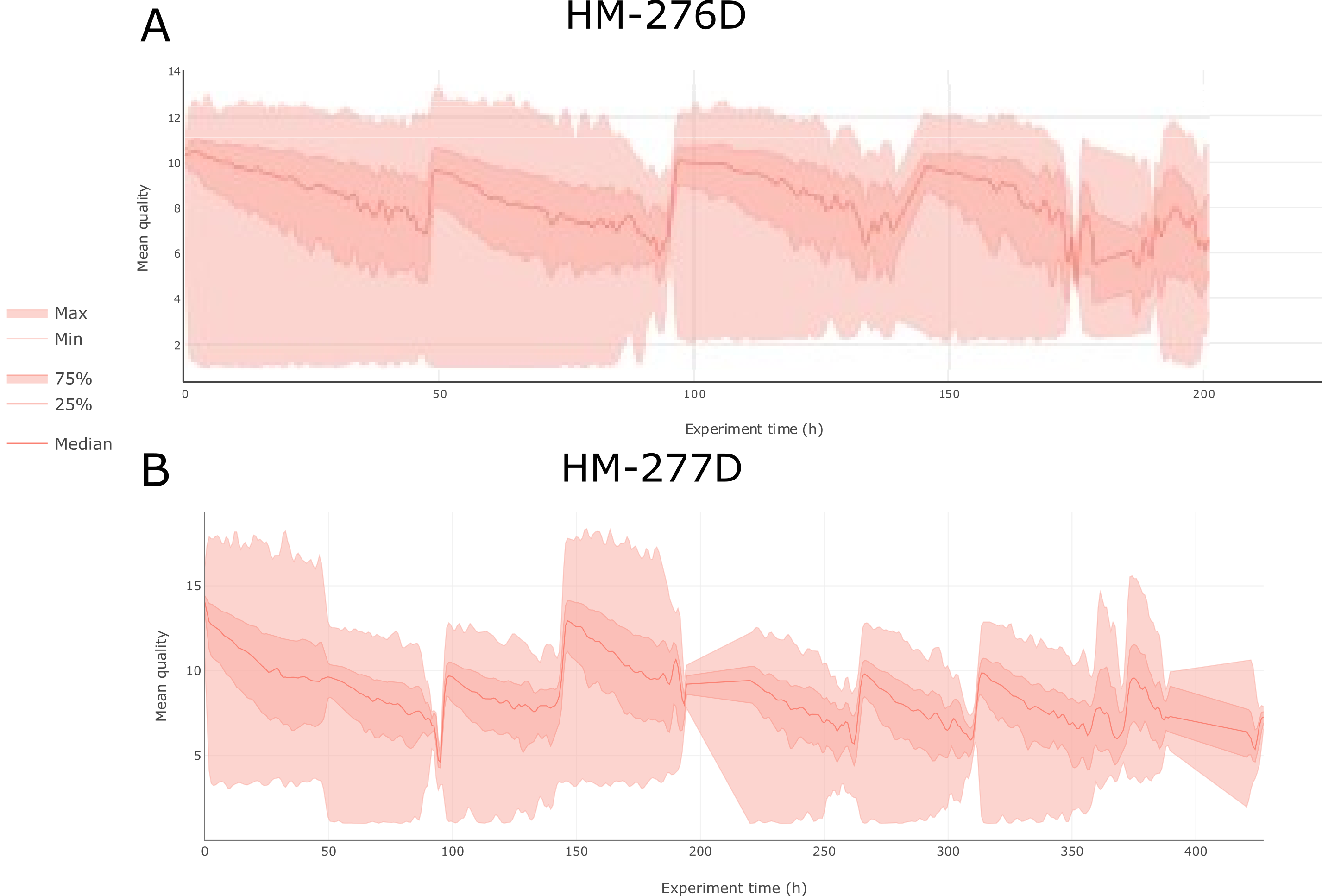

**Figure S4. Read quality over experiment of Nanopore sequencing data.** Mean read quality over experiment time for sequenced data sets, HM-276D **(A)** and HM-277D **(B)**, were summarized using PycoQC.

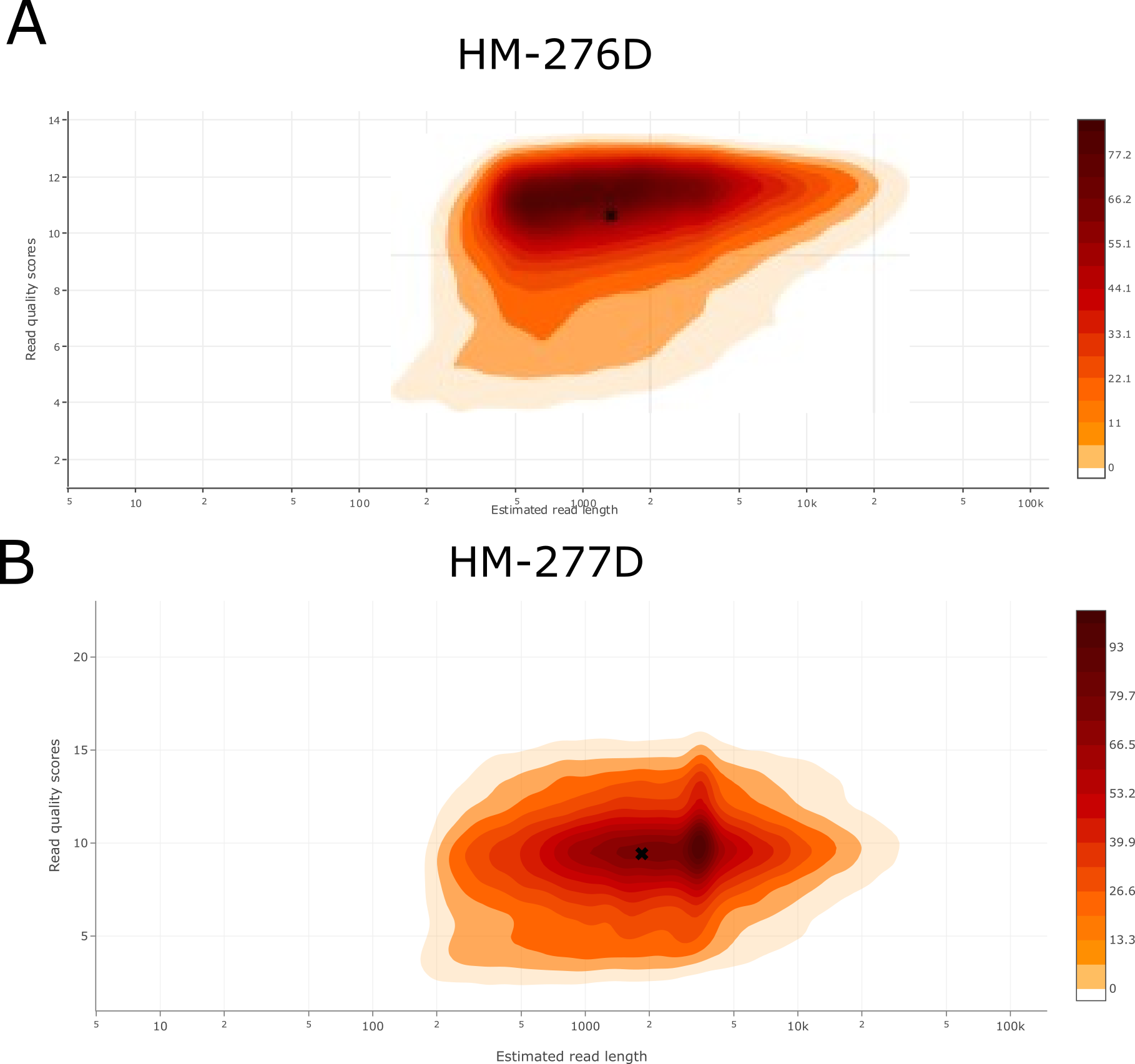

**Figure S5. Read quality score vs estimated read length.** Nanopore read distribution of read length and quality score for sequenced data sets, HM-276D **(A)** and HM-277D **(B)**, were summarized using PycoQC. Color indicates read density.

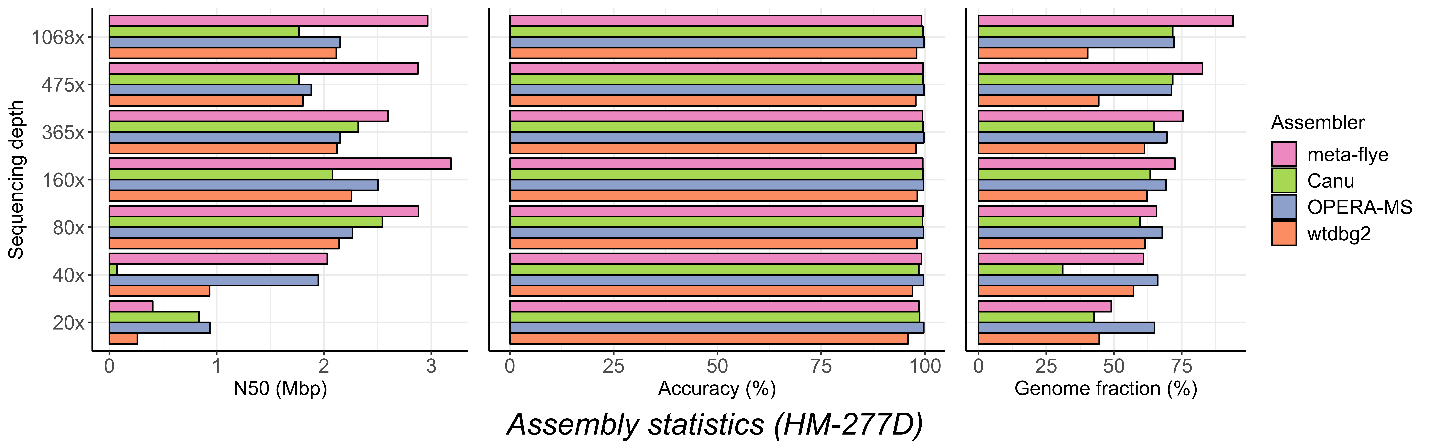

**Figure S6. Assembly performance on HM-277D data set.** Assembly statistics (N50 length, accuracy and genome fraction) of each assembler at different coverage depths based on HM-277D data set. Colors indicate results from different assemblers (Canu, OPERA-MS, wtdbg2, meta-flye). Assembly accuracy remains high compared to HM-276D, ranging around ~99% across tools. N50 lengths and genome fractions of all methods are substantially lower than the even community.

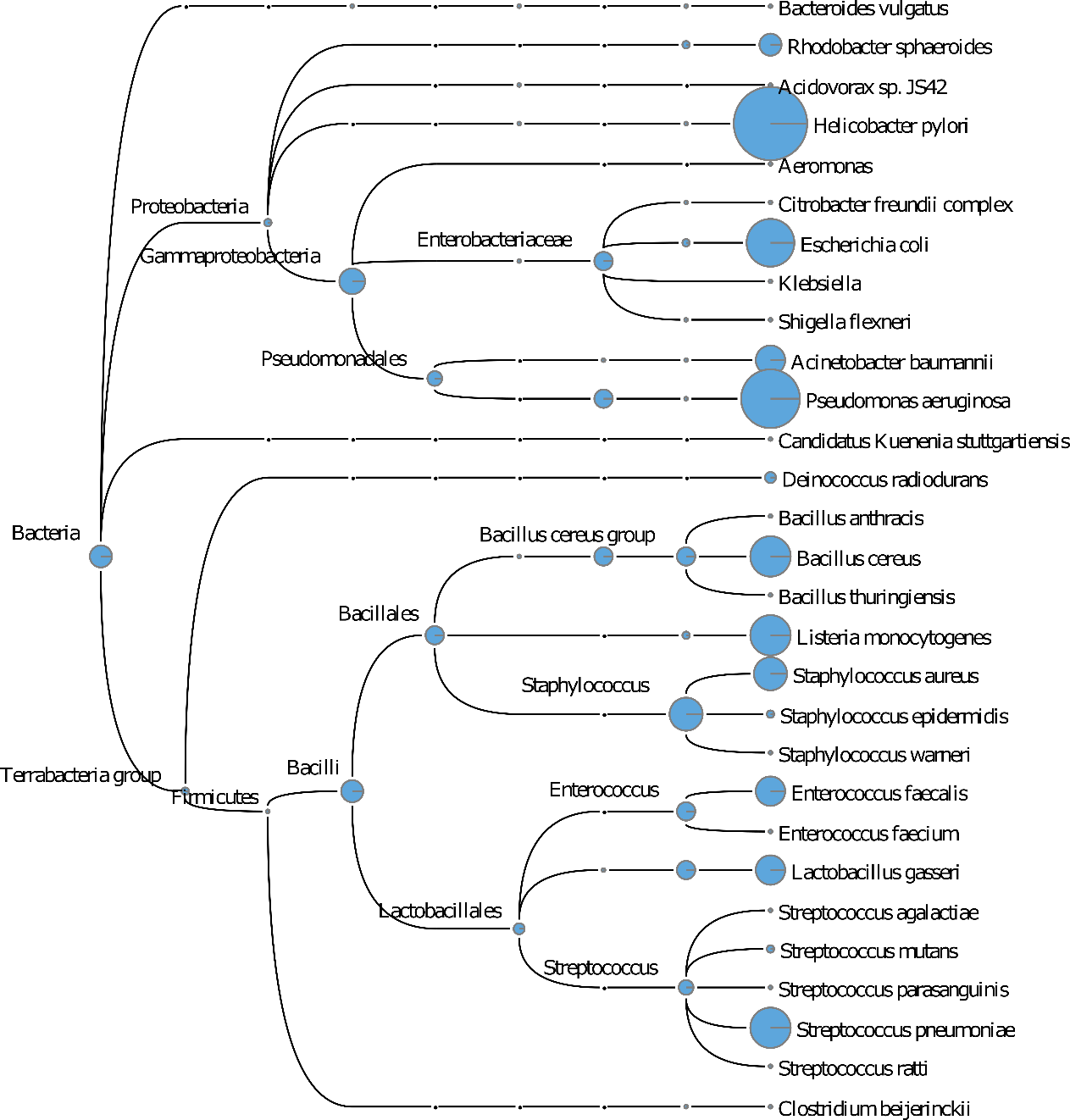

**Figure S7. Megan taxonomic tree assignment obtained from HM-276D PacBio sequenced data set.** HM-276D PacBio data set was subsampled to 160× depth of coverage. Each read was aligned against NCBI-nr protein reference data base, then binned and visualized using Megan-LR. Megan taxonomic tree showing bacteria taxa identified and their corresponding abundances across taxonomic rank. The radius of circle represents the number of reads assigned for each taxa.

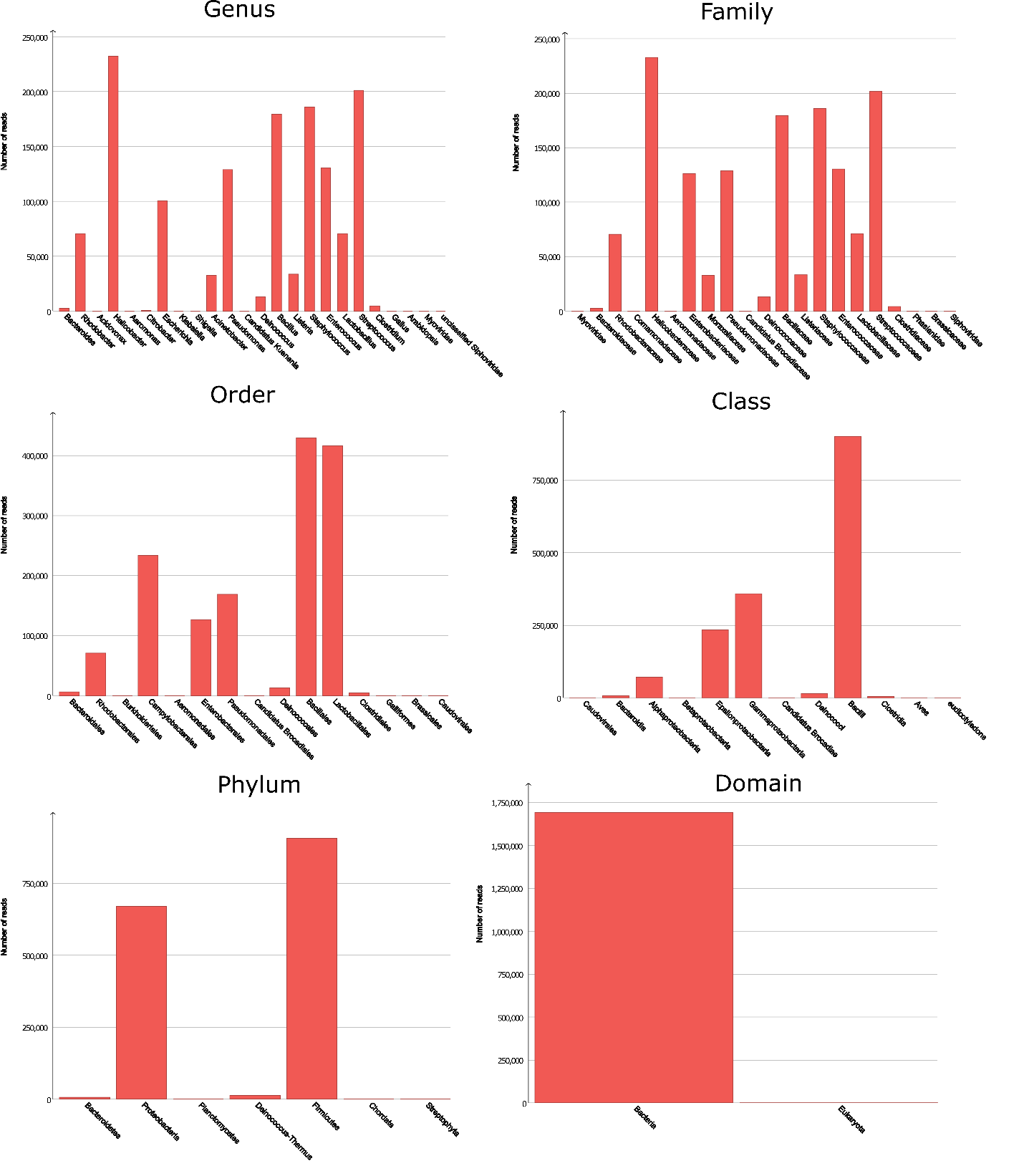

**Figure S8. Megan taxonomic read distribution at different ranks obtained from HM-276D Nanopore sequenced data set.** HM-276D Nanopore data set was subsampled to 160× depth of coverage. Each read was aligned against NCBI-nr protein reference data base, then binned and visualized using Megan-LR.

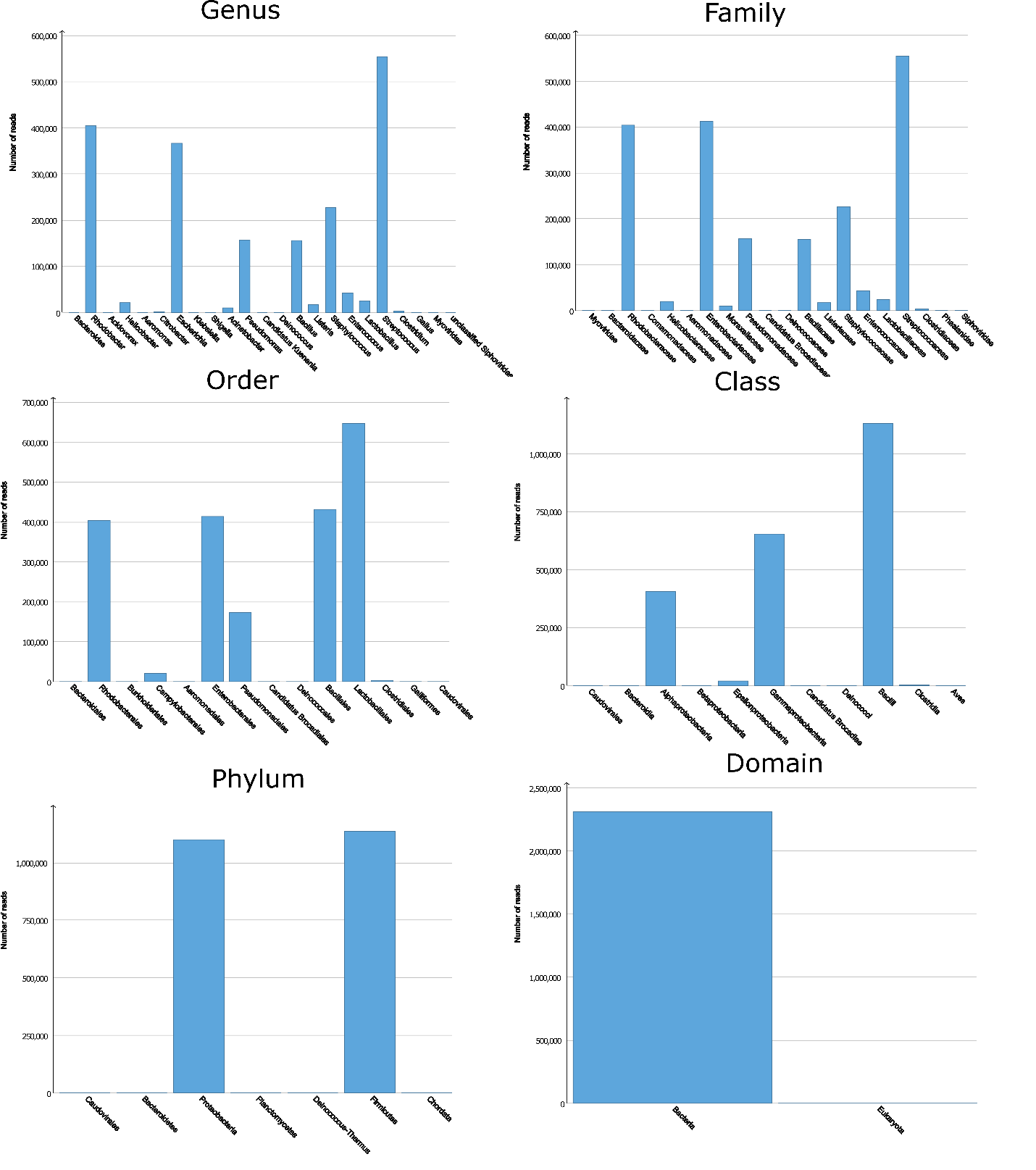

**Figure S9. Megan taxonomic read distribution at different ranks obtained from HM-277D Nanopore sequenced data set.** HM-277D Nanopore data set was subsampled to 160× depth of coverage. Each read was aligned against NCBI-nr protein reference data base, then binned and visualized using Megan-LR.

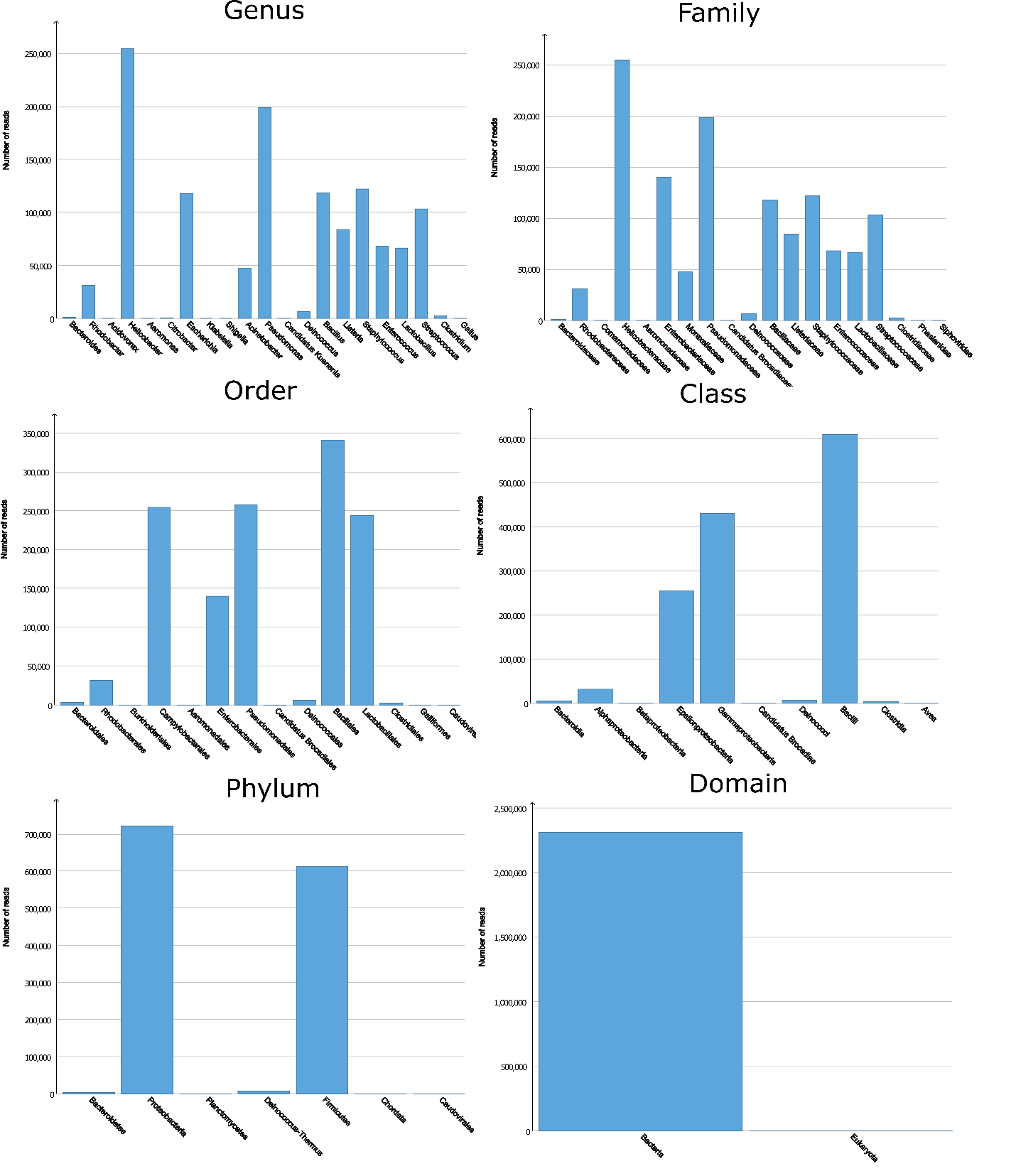

**Figure S10. Megan taxonomic read distribution at different ranks obtained from HM-276D PacBio sequenced data set.** HM-276D PacBio data set was subsampled to 160× depth of coverage. Each read was aligned against NCBI-nr protein reference data base, then binned and visualized using Megan-LR.

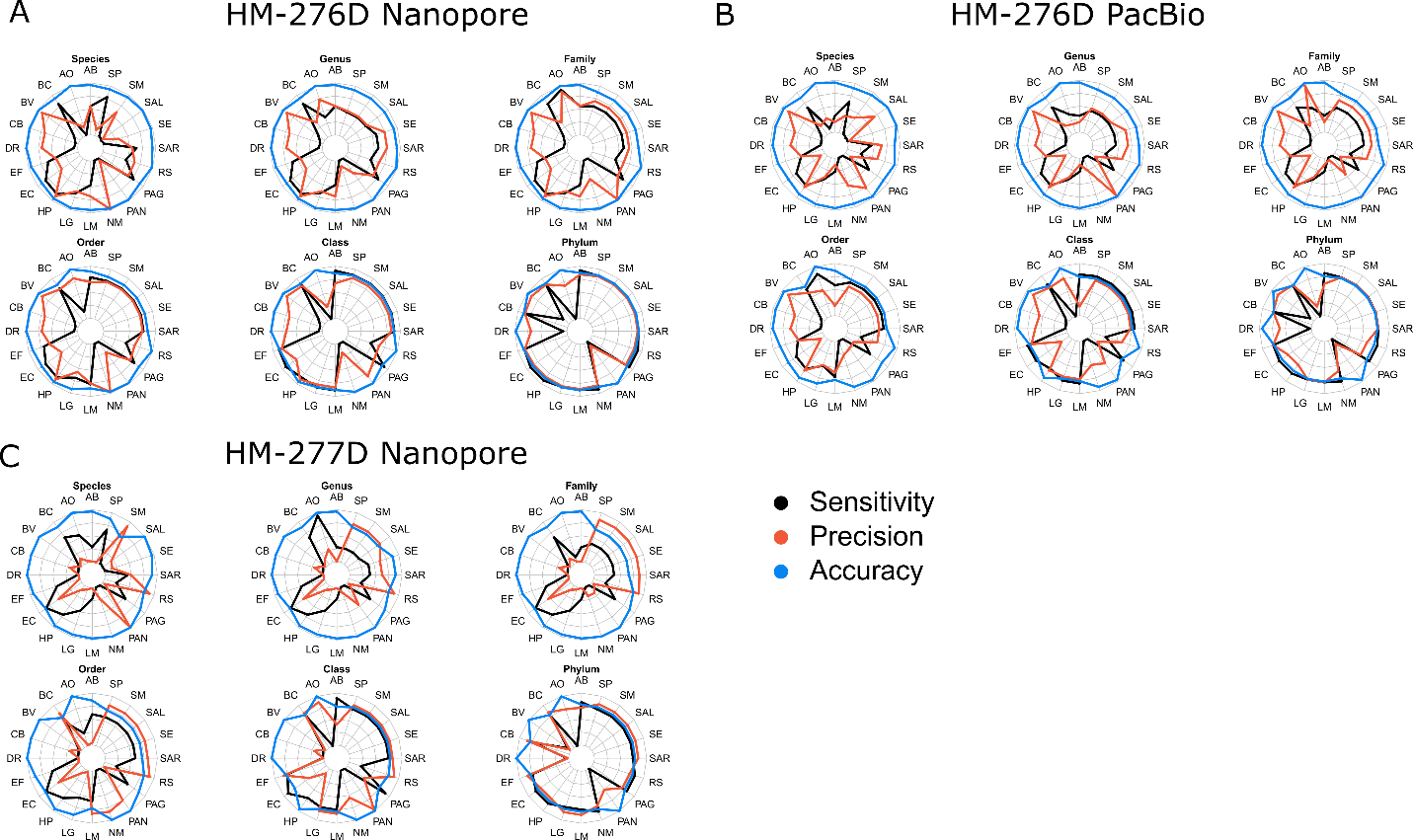

**Figure S11. Strain-specific read assignment performance comparison across sequencing technologies.** Read assignment accuracy statistics for each bacterial strain were summarized based on datasets: HM-276D Nanopore **(A)**, HM-276D PacBio **(B)** and HM-277D Nanopore **(C)** across ranks. Colors indicates different metrics: sensitivity, precision and accuracy. Taxon were accurately recovered above the family level. HM-276D Nanopore outperformed other two data sets. AB, *Acinetobacter baumannii*; AO, *Actinomyces odontolyticus*; BC, *Bacillus cereus*; BV, *Bacteroides vulgatus*; CB, *Clostridium beijerinckii*; DR, *Deinococcus radiodurans*; DF, *Enterococcus faecalis*; EC, *Escherichia coli*; HP, *Helicobacter pylori*; LG, *Lactobacillus gasseri*; LM, *Listeria monocytogenes*; NM, *Neisseria meningitides*; PAN, *Propionibacterium acnes*; PAG, *Pseudomonas aeruginosa*; RS, *Rhodobacter sphaeroides*; SAR, *Staphylococcus aureus*; SE, *Staphylococcus epidermidis*; SAL, *Streptococcus agalactiae*; SM, *Streptococcus mutans*; SP, *Streptococcus pneumonia*.

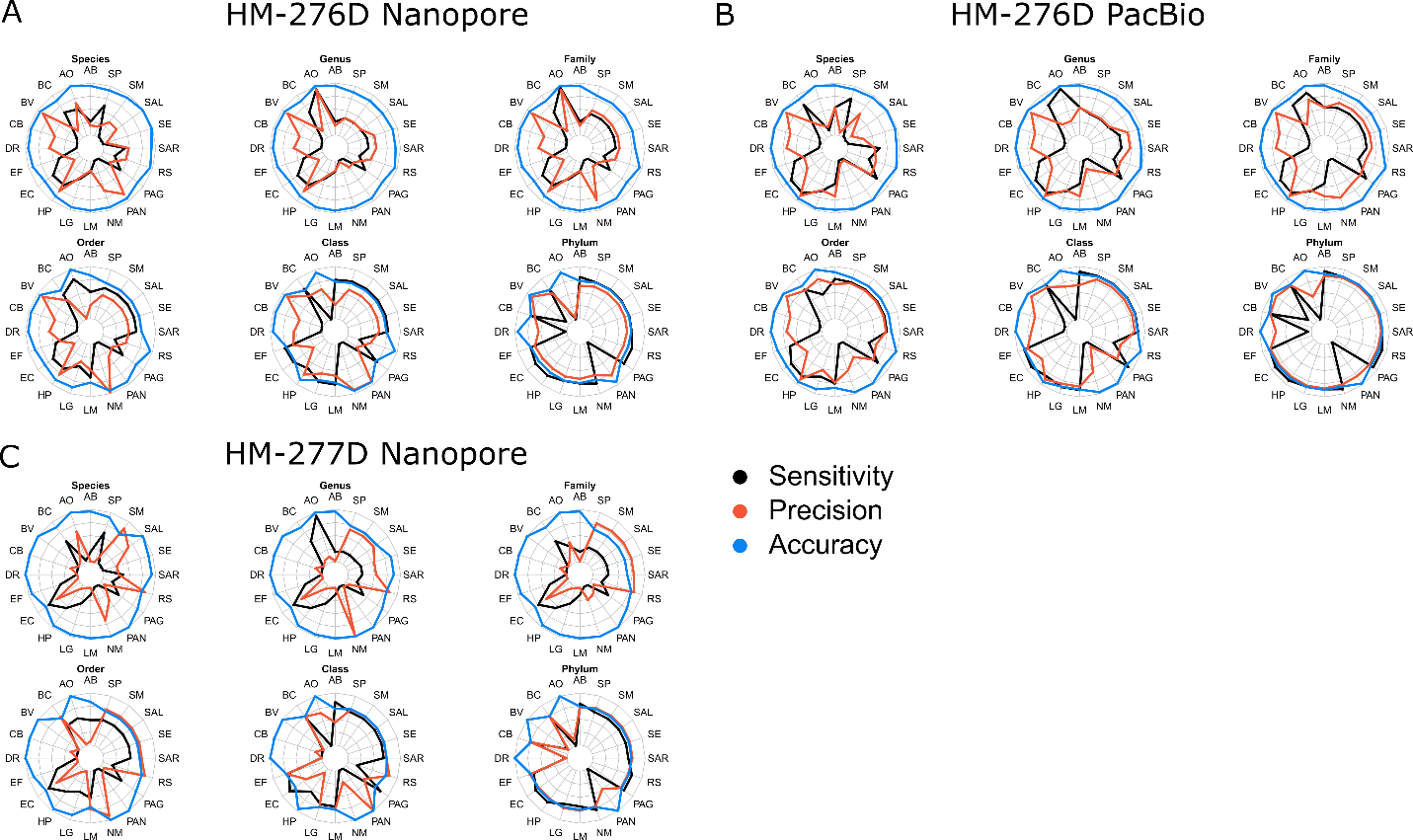

**Figure S12. Strain-specific base pair assignment performance comparison across sequencing technologies.** Read base assignment accuracy statistics for each bacterial strain were summarized based on datasets: HM-276D Nanopore **(A)**, HM-276D PacBio **(B)** and HM-277D Nanopore **(C)** across ranks. Colors indicates different metrics: sensitivity, precision and accuracy. PacBio performed better than Nanopore data above the family level because of lower error rate. AB, *Acinetobacter baumannii*; AO, *Actinomyces odontolyticus*; BC, *Bacillus cereus*; BV, *Bacteroides vulgatus*; CB, *Clostridium beijerinckii*; DR, *Deinococcus radiodurans*; DF, *Enterococcus faecalis*; EC, *Escherichia coli*; HP, *Helicobacter pylori*; LG, *Lactobacillus gasseri*; LM, *Listeria monocytogenes*; NM, *Neisseria meningitides*; PAN, *Propionibacterium acnes*; PAG, *Pseudomonas aeruginosa*; RS, *Rhodobacter sphaeroides*; SAR, *Staphylococcus aureus*; SE, *Staphylococcus epidermidis*; SAL, *Streptococcus agalactiae*; SM, *Streptococcus mutans*; SP, *Streptococcus pneumonia*.

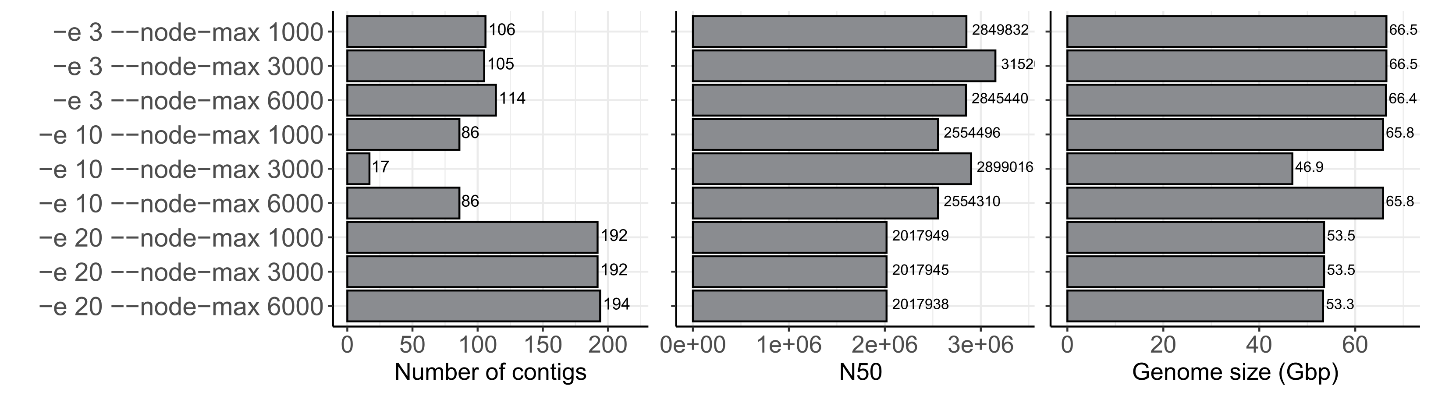

**Figure S13.** **Assembly results for wtdbg2 based on HM-276D data sets.** Barplots indicate assembly statistics (Number of contigs, N50 length and genome size). Each row represents a run of wtdbg2 with parameters “-e” and “--node-max”.

| **Tools** | **Depth** | **N50 length** | **NG50 length** | **NA50 length** | **NGA50 length** | **Accuracy (%)** | **Fraction (%)** | **# contigs** | **# long contigs** | **Longest contig** | **Genome size** | **CPU time (min)** |
| --- | --- | --- | --- | --- | --- | --- | --- | --- | --- | --- | --- | --- |
| **Canu** | **20x** | 717267 | 616530 | 450953 | 413727 | 98.5 | 96.8 | 298 | 254 | 2612567 | 65503873 | 98 |
| **Canu** | **40x** | 1987236 | 1975600 | 893966 | 893966 | 99.07 | 99.29 | 132 | 112 | 6286130 | 67676017 | 250 |
| **Canu** | **80x** | 2886059 | 2731942 | 1284219 | 1284219 | 99.24 | 99.86 | 62 | 57 | 6316623 | 68735511 | 678 |
| **Canu** | **160x** | 3901381 | 3901381 | 1302124 | 1506778 | 99.27 | 99.93 | 60 | 52 | 6299115 | 68879111 | 1537 |
| **Canu** | **365x** | 2983818 | 2983818 | 1219953 | 1219953 | 99.28 | 99.83 | 64 | 58 | 6292103 | 68964121 | 2651 |
| **Canu** | **480x** | 3911963 | 3911963 | 1400993 | 1400993 | 99.4 | 99.81 | 83 | 65 | 6359094 | 69425747 | 3858 |
| **OPERA-MS** | **20x** | 1122204 | 1122204 | 450323 | 489324 | 99.83 | 99.71 | 5117 | 201 | 6324007 | 67168904 | 61 |
| **OPERA-MS** | **40x** | 2657727 | 2657727 | 1210958 | 1210958 | 99.96 | 99.99 | 1695 | 81 | 5220208 | 67629371 | 102 |
| **OPERA-MS** | **80x** | 2835709 | 2732545 | 1189226 | 1189226 | 99.96 | 99.99 | 1921 | 74 | 4636570 | 67632885 | 186 |
| **OPERA-MS** | **160x** | 2933262 | 2792941 | 1273171 | 1273171 | 99.95 | 98.45 | 2347 | 65 | 6255842 | 66580943 | 382 |
| **OPERA-MS** | **365x** | 2938016 | 2938016 | 1298425 | 1298425 | 99.91 | 99.98 | 4734 | 64 | 6255878 | 67858470 | 856 |
| **OPERA-MS** | **480x** | 2938019 | 2938019 | 1213537 | 1298125 | 99.92 | 99.98 | 4732 | 63 | 6255756 | 67892051 | 1238 |
| **wtdbg2** | **20x** | 552100 | 415407 | 243026 | 208370 | 96.22 | 90.73 | 439 | 367 | 3542441 | 60910472 | 3 |
| **wtdbg2** | **40x** | 2106610 | 2057746 | 751004 | 751004 | 98.1 | 98.58 | 143 | 110 | 6265362 | 66286613 | 7 |
| **wtdbg2** | **80x** | 3152112 | 2920474 | 1331021 | 1331021 | 98.79 | 98.97 | 105 | 66 | 6229939 | 66511311 | 13 |
| **wtdbg2** | **160x** | 2910424 | 2910424 | 1286826 | 1286826 | 98.95 | 98.75 | 134 | 76 | 6215258 | 66615029 | 13 |
| **wtdbg2** | **365x** | 2706821 | 2706821 | 1278483 | 1265683 | 98.66 | 97.34 | 90 | 73 | 6251621 | 65544245 | 19 |
| **wtdbg2** | **480x** | 3168384 | 2922530 | 1201346 | 1184259 | 98.73 | 95.95 | 201 | 119 | 6210210 | 65977641 | 23 |
| **meta-flye** | **20x** | 1653589 | 1547909 | 558652 | 534341 | 98.96 | 98.76 | 223 | 206 | 5630982 | 66808399 | 57 |
| **meta-flye** | **40x** | 2725547 | 2653197 | 1209147 | 1209147 | 99.43 | 99.97 | 64 | 52 | 6274273 | 67627825 | 86 |
| **meta-flye** | **80x** | 2930772 | 2930772 | 1588493 | 1588493 | 99.52 | 99.99 | 59 | 43 | 6251934 | 67630110 | 140 |
| **meta-flye** | **160x** | 3888260 | 3180529 | 1622636 | 1622636 | 99.54 | 99.97 | 61 | 39 | 6252579 | 67595608 | 372 |
| **meta-flye** | **365x** | 3181836 | 2934283 | 1315358 | 1315358 | 99.62 | 99.98 | 88 | 44 | 6245780 | 67727067 | 603 |
| **meta-flye** | **480x** | 3181822 | 2934277 | 1718698 | 1718698 | 99.62 | 99.99 | 89 | 43 | 6245565 | 67700317 | 756 |

**Table S1. Comprehensive assembly statistics on HM-276D using Canu, OPERA-MS, wtdbg2 and meta-flye.**

| **Species** | **Abundance** | **RefSeq gene** | | **16S rRNA gene** | | | **Protein coding gene** | |
| --- | --- | --- | --- | --- | --- | --- | --- | --- |
|  |  | average coverage(#bases) | Significantly detected gene | average coverage(#bases) | | Significantly detected gene | average coverage(#bases) | Significantly detected gene |
| **Acinetobacter baumannii** | 0.18% | 9.83 | 94 | 9.50 | 6 | | 9.86 | 3,817 |
| **Actinomyces odontolyticus** | 0.01% | 4.27 | 56 | 3.10 | 2 | | 4.65 | 1,999 |
| **Bacillus cereus** | 1.22% | 100.51 | 138 | 94.04 | 12 | | 102.33 | 5,675 |
| **Bacteroides vulgatus** | 0.02% | 2.32 | 65 | 1.77 | 4 | | 2.39 | 3,067 |
| **Clostridium beijerinckii** | 1.43% | 96.40 | 143 | 78.49 | 14 | | 97.42 | 5,149 |
| **Deinococcus radiodurans** | 0.03% | 4.94 | 57 | 5.19 | 3 | | 4.86 | 3,060 |
| **Enterococcus faecalis** | 0.01% | 2.76 | 53 | 3.81 | 2 | | 3.37 | 2,497 |
| **Escherichia coli** | 15.75% | 1,032.93 | 179 | 1,003.79 | 7 | | 1,060.46 | 4,341 |
| **Helicobacter pylori** | 0.07% | 113.13 | 43 | 117.15 | 2 | | 114.16 | 1,444 |
| **Lactobacillus gasseri** | 0.03% | 27.95 | 96 | 24.06 | 6 | | 28.97 | 1,783 |
| **Listeria monocytogenes** | 0.07% | 10.74 | 184 | 8.92 | 6 | | 11.42 | 2,864 |
| **Neisseria meningitides** | 0.07% | 42.67 | 71 | 28.53 | 4 | | 47.85 | 1,926 |
| **Propionibacterium acnes** | 0.11% | 41.60 | 58 | 38.75 | 3 | | 43.02 | 2,506 |
| **Pseudomonas aeruginosa** | 5.01% | 141.55 | 105 | 160.86 | 4 | | 137.90 | 5,572 |
| **Rhodobacter sphaeroides** | 64.44% | 2,219.40 | 67 | 1,993.22 | 3 | | 2,438.52 | 4,279 |
| **Staphylococcus aureus** | 0.83% | 323.26 | 79 | 289.00 | 5 | | 404.68 | 2,982 |
| **Staphylococcus epidermidis** | 6.52% | 976.37 | 76 | 1,117.10 | 5 | | 1,002.43 | 2,472 |
| **Streptococcus agalactiae** | 0.03% | 72.99 | 101 | 70.16 | 7 | | 75.54 | 2,127 |
| **Streptococcus mutans** | 4.15% | 4,207.60 | 80 | 3,598.02 | 5 | | 3,818.93 | 1,953 |
| **Streptococcus pneumoniae** | 0.01% | 1.91 | 58 | 1.30 | 2 | | 2.39 | 1,868 |

**Table S2. Species-specific gene coverage summary of HM-277D data set.** Gene coverage statistics were summarized for 3 different gene sets: all Refseq genes, 16S rRNA genes and protein coding genes. Average coverage = number of bases mapped to the exonic region / length of exonic region. Gene is noted as significantly detected when 50% exonic region is covered by at least 1 read and average coverage > 1.

### ****Transparent Methods****

#### Oxford nanopore sequencing of HM-276D and HM-277D

DNA samples of HM-276D and HM-277D were ordered from BEI Resources. Concentration of DNA was assessed using the dsDNA HS assay on a Qubit fluorometer (Thermo Fisher).

For library preparation, 1.0 µg DNA was used as the input DNA of each library. The library was prepared using the ligation sequencing protocol (SQK-LSK109) from ONT. Concretely, end repair, dA-tailing and DNA repair was performed using NEBNext Ultra II End Repair/dA-tailing Module (catalog No. E7546) and NEBNext FFPE Repair Mix (M6630). In all, 3.5 μl Ultra II End-prep reaction buffer, 3 μl Ultra II End-prep enzyme mix, 3.5 μl NEBNext FFPE DNA Repair Buffer and 2 μl NEBNext FFPE DNA Repair Mix were added to the input DNA. The total volume was adjusted to 60 µl by adding nuclease-free water (NFW). The mixture was incubated at 20 °C for 5 min and 65 °C for 5 min. A 1 × volume (60 µl) AMPure XP clean-up was performed and the DNA was eluted in 61 µl NFW. One microliter of the eluted dA-tailed DNA was quantified using the Qubit fluorometer. A total of 0.7 µg DNA should be retained if the process is successful.

Adaptor ligation was performed using the following steps. Five microliter Adaptor Mix (ONT, SQK-LSK109 Kit), 25 μl Ligation Buffer (ONT, SQK-LSK109 Kit) and 10 μl NEBNext Quick T4 DNA Ligase (NEB, catalog No. E6056) were added to the 60 µl dA-tailed DNA from the previous step. The mixture was incubated at room temperature for 10 min. The adaptor-ligated DNA was cleaned up using 40 µl AMPure XP beads. The mixture of DNA and AMPure XP beads was incubated for 5 min at room temperature and the pellet was washed twice by 250 μl Long Fragment Buffer (ONT, SQK-LSK109). The purified-ligated DNA was resuspended in 15 µl Elution Buffer (ONT, SQK-LSK109). A 1-µl aliquot was quantified by fluorometry (Qubit) to ensure ≥ 400 ng DNA was retained. The final library was prepared by mixing 37.5 μl Sequencing Buffer (ONT, SQK-LSK109), 25.5 μl Loading Beads (ONT, SQK-LSK109), and 12 µl purified-ligated DNA. The library was loaded to R9.4 flow cells (FLO-MIN106, ONT) according to the manufacturer’s guidelines. GridION sequencing was performed using default settings for the R9.4 flow cell and SQK-LSK109 library preparation kit. The sequencing was controlled and monitored using the MinKNOW software developed by ONT.

#### Metagenome assembly

Genome assemblies of the 20-mixed bacteria from HM-276D and MH-277D mock communities were conducted using 4 existing assemblers based on generated long-read sequencing reads. These 4 dedicated long-read assemblers we used are wtdbg2 (v2.4), OPERA-MS, Canu (v1.8) and meta-flye, where OPERA-MS and meta-flye are designed to be capable to handle metagenome while wdtbg2 and Canu are for broadly application. To evaluate the impact of coverage depth in genome assembly, in addition to 525× (HM-276D) and 1068× (HM-277D), we subsampled 5 data sets with 365×, 160×, 80×, 40× and 20× coverages for these two mock communities. In addition to long-read data, OPERA-MS requires short reads to improve the assembly accuracy. Hence, we downloaded Illumina sequenced HM-276D(Jones et al., 2015) and HM-277D data sets(Kuleshov et al., 2016). Similarly, these short-read data were also subsampled with depths 160×, 80×, 40× and 20×, which were provided to OPERA-MS in corresponding data set analysis. We also analyzed a PacBio data set(Lee et al., 2014) of HM-276D sample using wtdbg2, OPERA-MS, Canu and meta-flye to compare assembly performance across sequencing technologies. For comparison fairness, we applied consistent configuration settings for each tool across different coverage depths. For wtdbg2, we first tuned parameters on “-e”, “--node-max”. Assemblies were conducted under different parameter values (-e: 3, 10, 20) (--node-max: 1000, 3000, 6000). Based on the genome contiguity and completeness results in **Supplementary Fig 13**, we specified parameter “-e 3 –node-max” for wtdbg2. For other tools, we set estimated genome size as 70M, where the parameters are “genomeSize=70M useGrid=True” for Canu, and “CONTIG_LEN_THR 500, CONTIG_EDGE_LEN 80, CONTIG_WINDOW_LEN 340, KMER_SIZE 60, LONG_READ_MAPPER minimap2” for OPERA-MS, “-t 40 -g 70m -o ./ --meta” for meta-flye. 40 contig output files were obtained (2 mock community samples, 6 depths of coverage, 4 assembly tools) for further evaluation.

#### Metagenome assembly evaluation

Assembled genomes produced by each tool based on different samples and coverage depths were evaluated with metrics related to contiguity, genome completeness and accuracy. To assess the assembly contiguity, we first used our script to calculate the widely-used statistic N50, which is the shortest contig needed to cover at least 50% of the assembly. In addition, other related statistics, such as number of contigs, number of long contigs (>10kb), longest contigs and total assembly size, were collected from the FASTA output file of each assembler. Furthermore, we summarized NG50 for each method by replacing the assembly size with estimated genome size. This quantity represents the shortest contig needed to cover 50% of the genome. Instead of contigs, we also evaluated the performance based on aligned blocks for each method by using QUAST(Gurevich et al., 2013) to calculate NA50 and NGA50, which represent shortest aligned block to cover 50% of the assembly and genome respectively. Based on these metrics, the contiguity of assemblies was comprehensively evaluated. Next, we downloaded the reference genome FASTA files of all 20 bacteria from NCBI database to measure the concordance between the references and assemblies. First, assemblies were mapped to the reference genomes using Mummer v3.23 with parameters “-maxmatch -c 100 -p nucmer”. Then, by comparing all contigs mapped onto the reference using dandiff, assembly accuracy was calculated using 1-to-1 alignment identity, which is the correctly matched base-pair percentage of contigs uniquely mapped to the reference genome (1-mismatch%). In addition, to assess the assembly completeness, we calculated the percentage of genome covered by the contigs. In real case, instead of evenly mixed in HM-276D mock community, bacterial strains are non-uniformly distributed, where some are likely to share extremely low abundance. Therefore, we evaluated the impact of the genomic DNA abundance on genome assembly. For the unevenly mixed HM-277D mock community samples, we calculated the abundance for each bacterial strain by normalizing the concentration with related reference genome size. The relationship between abundances and assessment metrics was displayed using scatter plots. For each plot, linearity was measured based on Spearman correlation using R v3.3.3.

#### Taxonomic binning analysis

Taxon bins of the 20-mixed bacteria from two mock communities were recovered using taxonomic binner Megan-LR(Huson et al., 2018) with 3 long-read sequencing data sets: HM-276D (Nanopore, PacBio) and HM-277D (Nanopore) at 160× depth of coverage. We first aligned all reads against NCBI-nr protein reference database using LAST with parameters “-P 100 -F15”. Next, output MAF files were converted to DAA format in smaller size. Then, we meganized the DAA files using MEGAN(Huson et al., 2016), which allows us to interactively visualize and explore these taxonomic results. To evaluate the taxonomic binning performance, we first counted the number of reads and bases which were correctly assigned to each taxon from the mock microbial community. We determined the metrics (precision, sensitivity, true positive rate and false positive rate). Precision and sensitivity assess how accuracy each read is classified across different sequencing technologies. Precision is the percentage of reads assigned correctly to the corresponding taxa out of all reads. Sensitivity is the percentage of correct reads out of reads assigned to the particular taxa. Next, we use true positive rate (TPR) and false discover rate (FDR) to assess the accuracy in taxonomic detection across sequencing technologies. TPR is the percentage of correctly detected taxon out of known taxon from the microbial community. FDR is the percentage of correctly detected taxon out of all detected taxon. All metrics are defined at each taxonomic rank.

### ****Supplemental References****

GUREVICH, A., SAVELIEV, V., VYAHHI, N. & TESLER, G. 2013. QUAST: quality assessment tool for genome assemblies. *Bioinformatics,* 29**,** 1072-5.

HUSON, D. H., ALBRECHT, B., BAGCI, C., BESSARAB, I., GORSKA, A., JOLIC, D. & WILLIAMS, R. B. H. 2018. MEGAN-LR: new algorithms allow accurate binning and easy interactive exploration of metagenomic long reads and contigs. *Biol Direct,* 13**,** 6.

HUSON, D. H., BEIER, S., FLADE, I., GORSKA, A., EL-HADIDI, M., MITRA, S., RUSCHEWEYH, H. J. & TAPPU, R. 2016. MEGAN Community Edition - Interactive Exploration and Analysis of Large-Scale Microbiome Sequencing Data. *PLoS Comput Biol,* 12**,** e1004957.

JONES, M. B., HIGHLANDER, S. K., ANDERSON, E. L., LI, W., DAYRIT, M., KLITGORD, N., FABANI, M. M., SEGURITAN, V., GREEN, J., PRIDE, D. T., YOOSEPH, S., BIGGS, W., NELSON, K. E. & VENTER, J. C. 2015. Library preparation methodology can influence genomic and functional predictions in human microbiome research. *Proc Natl Acad Sci U S A,* 112**,** 14024-9.

KULESHOV, V., JIANG, C., ZHOU, W., JAHANBANI, F., BATZOGLOU, S. & SNYDER, M. 2016. Synthetic long-read sequencing reveals intraspecies diversity in the human microbiome. *Nat Biotechnol,* 34**,** 64-9.

LEE, C. H., BOWMAN, B. & HALL, R. Developments in PacBio® metagenome sequencing: Shotgun whole genomes and full-length 16S. International Plant and Animal Genome Conference Asia, 2014.
